# Structure and lipid-mediated remodelling mechanism of the Na^+^/H^+^ exchanger NHA2

**DOI:** 10.1101/2021.07.22.453398

**Authors:** Rei Matsuoka, Roman Fudim, Sukkyeong Jung, Chenou Zhang, Andre Bazzone, Yurie Chatzikyriakidou, Norimichi Nomura, So Iwata, Laura Orellana, Oliver Beckstein, David Drew

## Abstract

Na^+^/H^+^ exchangers catalyse an ion-exchange activity that is carried out in most, if not all cells. SLC9B2, also known as NHA2, correlates with the long-sought after sodium/lithium (Na^+^/Li^+^) exchanger linked to the pathogenesis of diabetes mellitus and essential hypertension in humans. Despite its functional importance, structural information and the molecular basis of its ion-exchange mechanism have been lacking. Here, we report the cryo EM structures of *bison* NHA2 in detergent and in nanodiscs at 3.0 and 3.5 Å resolution, respectively. NHA2 shares closest structural similarity to the bacterial electrogenic Na^+^/H^+^ antiporter NapA, rather than other mammalian SLC9A members. Nevertheless, SSM-based electrophysiology results with NHA2 show the catalysis of electroneutral rather than electrogenic ion exchange, and the ion-binding site is quite distinctive, with a tryptophan-arginine-glutamate triad separated from the well-established ion-binding aspartates. These triad residues fine-tune ion binding specificity, as demonstrated by a salt-bridge swap mutant that converts NHA2 into a Li^+^-specific transporter. Strikingly, an additional N-terminal helix in NHA2 establishes a unique homodimer with a large ∼ 25 Å intracellular gap between protomers. In the presence of phosphatidylinositol lipids, the N-terminal helix rearranges and closes this gap. We confirm that dimerization of NHA2 is required for activity *in vivo*, and propose that the N-terminal helix has evolved as a lipid-mediated remodelling switch for regulation of transport activity.

---

Intracellular salt, pH and cell volume must be tightly regulated for cell survival^1^. The transmembrane exchange of H^+^ for Na^+^(Li^+^) ions by the action of Na^+^/H^+^ exchangers is central in this homeostatic process^1–3^. In bacteria, exchange activity also generate a Na^+^ gradient used for energizing other Na^+^-driven solute transporters^4^. In mammals, there are 13 distinct NHE orthologues that belong to the Cation:Proton Antiporter (CPA) superfamily: NHE1-9 also known as SLC9A1-9, NHA1-2 also known as SLC9B1-2, and sperm specific NHE also known as SLC9C ^2, 3, 5^. The CPA superfamily can be separated into CPA1 and CPA2 clades (http://www.tcdb.org/). Members of the CPA1 group are made up of the NHE1-9 family of electroneutral Na^+^(K^+^)/H^+^ exchangers, which are well-known for their important roles in human physiology, disease and drug targeting ^2, 6–8^. Recently, the first structures of mammalian NHE1^9^ and NHE9^10^ were reported, confirming the overall similarity to previously determined bacterial Na^+^/H^+^ antiporter structures that operate by an “elevator” alternating-access mechanism^10^. In contrast to the NHEs, the mammalian NHA1 and NHA2 belong to the CPA2 clade ^5, 11, 12^ and have greater sequence similarity to CPA2 members of bacterial origin ^12^ ^12^ (Fig. 1a). Pairwise alignment between NHA2 and bacterial electrogenic Na^+^/H^+^ antiporters shows that NHA2 harbors the conserved ion-binding aspartate residues that make up the well-described “DD-motif”^13, 14^, wherein the proton-motive-force energizes Na^+^ export. Consistently, it was observed that NHA2 co-localizes with V-ATPase^15, 16^ wherein cation extrusion is driven by an inwardly-directed proton gradient^17^, which is opposite to NHEs that are driven by Na^+^-gradients ^2, 3^ (Fig. 1b). This physiological direction, however, agrees with the observation that in flies, NHA2 expression specifically protected against sodium salt stress^16^. Based on tissue expression, genome location and phloretin sensitivity, *human* NHA2 was proposed to be the candidate gene for the Na^+^(Li^+^) countertransport activity linked to the development of essential hypertension and diabetes in humans^12, 17–19^. Consistently, NHA2 aids in sodium reabsorption in the kidney as a critical component of the WNK4-NCC pathway in the regulation of blood pressure homeostasis^20^. Furthermore, *in vitro* and *in vivo* studies show NHA2 as a contributor to insulin secretion in the β-cell^21^. NHA2 can be localized to both the plasma membrane in specialized cells and intracellular organelles including endosomes, lysosomes, and synaptic-like microvesicles^11, 15^. Here, we aimed to establish the structure and molecular-based mechanism of NHA2.

**Fig. 1.**
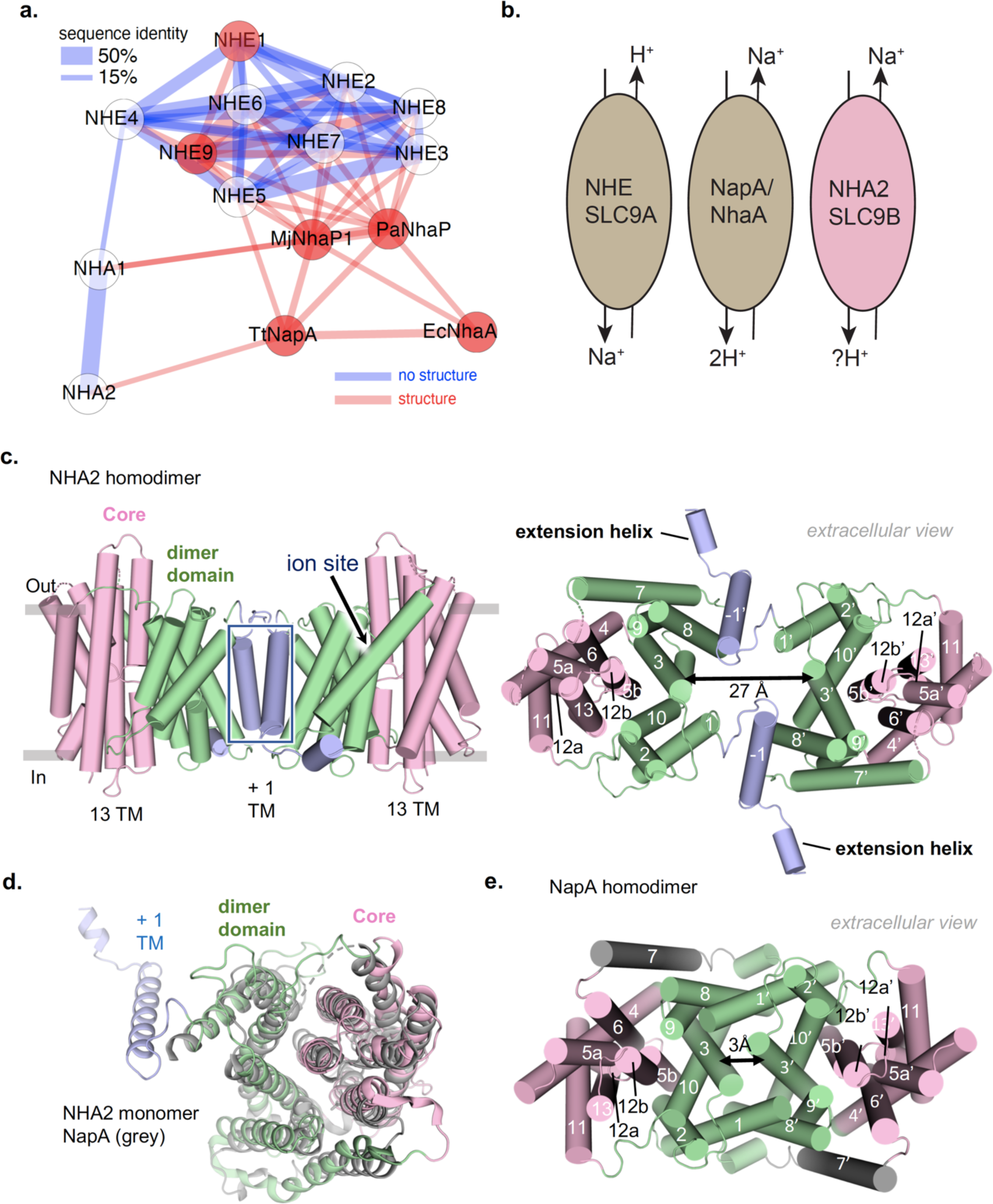
The cryo EM structure of NHA2 reveals a domain-swapped homodimer. **a**. Phylogenetic tree of canonical *human* NHE1-9 (SLC9A1-9) cluster compared to *human* NHA1 and NHA2 (SLC9B1-2) and bacterial members NapA (*Thermus thermophilus*), NhaP1 (*Methanococcus janashi*), NhaP (*Pyrococcus abyssi*) and NhaA (*Escherichia coli*). **b**. Schematic of the electroneutral NHE (SLC9A) CPA1 members that are driven by an inwardly directed Na^+^-gradient (left, brown)) as compared to electrogenic CPA2 Na^+^/H^+^ antiporters NapA and NhaA (middle, brown) that are driven by the proton-motive-force. The CPA2 member NHA2 is thought to be also driven by an inwardly-directed H^+^-gradient, yet activity is thought to be electroneutral. **c**. Cartoon representation of dimeric NHA2 from the side (left) and top view (right). Ion translocation 6-TM domain (transport) is colored in pink, dimerization domain (green) and the N-terminal domain-swapped transmembrane helix (TM–1) in blue. TM numbers are depicted in white. **d**. Cartoon representation of 14-TM NHA2 monomer from the extracellular side and coloured as in c. that has been superimposed onto to the 13-TM outward-facing structure of NapA (pdb: 4bwz) in grey. **e**. Cartoon representation of the NapA homodimer that has been coloured as NHA2 in c. to highlight that, in the absence of the additional N-terminal helix TM –1, an extensive and more compact oligomer is formed.

*Bison* NHA2 was selected for structural studies as it was more detergent stable in *S. cerevisiae* membranes than *human* NHA2 and several other mammalian homologues investigated (see Methods). The *bison* NHA2 structural construct is referred to as NHA2*Δ*N as it missing a 69 amino acid N-terminal tail deletion that is poorly conserved and was removed to improve expression and reduce predicted disorder (Methods, Supplementary Fig. 1); the shortened *bison* NHA2*Δ*N sequence shares 97% sequence identity to human NHA2*Δ*N (Supplementary Fig. 2, Extended Fig .1). Previously, it has been shown that only functional *human* NHA2 can rescue growth in the salt-sensitive *S. cerevisiae* Ab11c strain^12, 22^, which lacks the main Na^+^- and K^+^-extrusion systems. We confirmed that *bison* NHA2*Δ*N expressed in the Ab11c strain and complemented growth under Li^+^/Na^+^-salt stress conditions to at least the same extent as the full-length construct or *human* NHA2*Δ*N (Extended Fig. 2a-b, Supplementary Fig. 1a-b, Supplementary Fig. 3a). In contrast, poor *S. cerevisiae* growth was apparent under Na^+^/Li^+^-salt stress conditions for either non-induced cells or the double ion-binding aspartate mutant of *bison* NHA2*Δ*N (Asp278Cys-Asp279Cys), previously shown to abolish *human* NHA2 activity^23^ (Extended Fig. 2a-b, Supplementary Fig. 1a-b, Supplementary Fig. 3a). *Bison* NHA2*Δ*N was subsequently optimized for grid preparation, cryo-EM data acquisition, and structural determination at an active pH of 8.0 (Extended Data Fig 3 and Methods). Single-particle analysis (SPA) produced 2D class averages corresponding to predominantly side views of the NHA2*Δ*N homodimer. To improve 3D reconstruction, detergent subtraction and non-uniform refinement was performed in cryoSPARC (v.2.14.2). The final cryo-EM map was reconstructed to 3.0 Å, according to the gold-standard Fourier shell correlation (FSC) 0.143 criterion, which used ∼25% of the total collected particles (Table 1, Extended Data Fig. 3, Extended Data Fig. 4). Overall, the *bison* NHA2*Δ*N structure is well resolved, but due to inherent flexibility the connecting TM6 -TM7 loop could not be modelled (Methods).

**Fig. 2.**
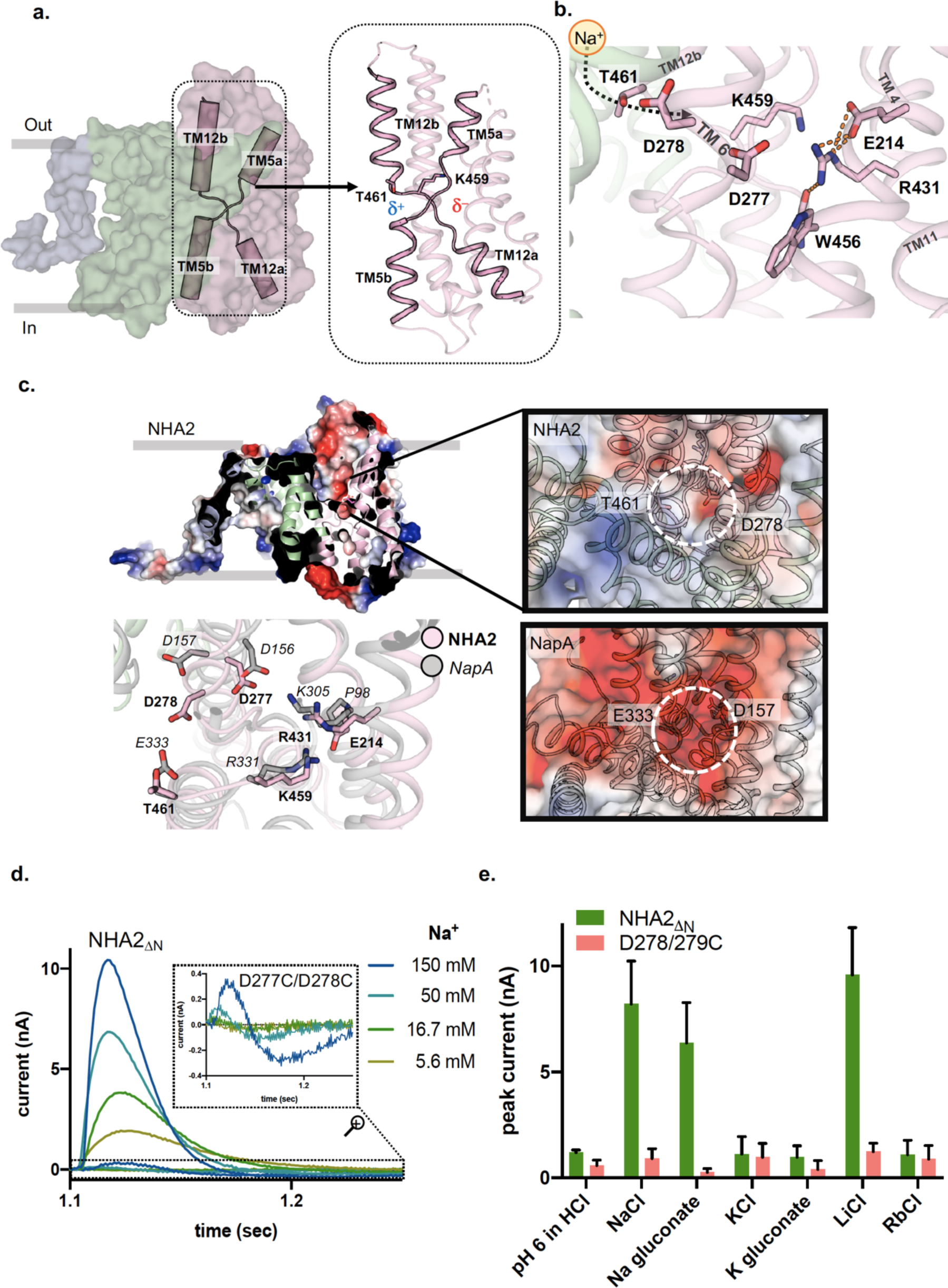
NHA2 ion-binding site and its electroneutral activity. **a.** Surface representation of NHA2 with the 6-TM core domain (pink), dimerization domain (green) and the N-terminal domain-swapped transmembrane helix TM –1 (blue). The crossover of half helices TM5a-b and TM12a-b (cartoon) are unique to the NhaA- fold and the half-helical dipoles that they create are highlighted. In NHA2, a lysine residue (K459 in stick form) is well-positioned to neutralise the negatively-charged half- helical dipoles, but the positively-charged dipoles lack a negatively-charged residue that is conserved in all other Na^+^/H^+^ antiporter structures^13^. Instead, the polar residue (T461 in stick form) that is conserved in all NHA2 members seems to be enough for stability (see ED Fig. 1). **b**. The ion-binding site of NHA2 has the two aspartates seen in electrogenic Na^+^/H^+^ antiporters (D278 and D277) and a NHA2-specific salt-bridge between R431 and E214 that is indirectly connected to W456 that are shown in stick- form. Dashed line represents hydrogen-bonding. **c**. *above left*: Cartoon representation of NHA2 with the electrostatic surface representation through the ion-binding site of one monomer (coloured blue to red, for positive to negative charges, respectively). The strictly conserved ion-binding residue Asp278 is labelled and shown in pink as sticks. *above right*: The outward open ion-binding cavity of NHA2 is encircled and highlighted. *below left*: The ion-binding site comparison between NHA2 (pink) and electrogenic NapA (grey), highlighting the differences in E214 (P98 in NapA) and T461 (E333) in NapA. *below right*: The outward open ion-binding cavity is comparatively more negatively-charged in NapA than NHA2 shown above. **d**. Transient currents recorded on *bison* NHA2 *Δ*N proteoliposomes under symmetrical pH 7.5 and increasing Na^+^ concentration jumps as shown; inset shows a zoomed in responses to *bison* NHA2 *Δ*N wherein the ion-binding aspartates were substituted to cysteine (D277C-D278C). **e**. peak current averages in response to different concentration jumps. The first bars show peak currents obtained by pH jumps from pH 7.0 to pH 6.0 titrated with HCl. The following bars show peak currents upon exchange of 150 mM choline chloride with 150 mM of the given salt at pH 7.5. Averages of recordings performed on five individual sensors are shown.

**Fig. 3.**
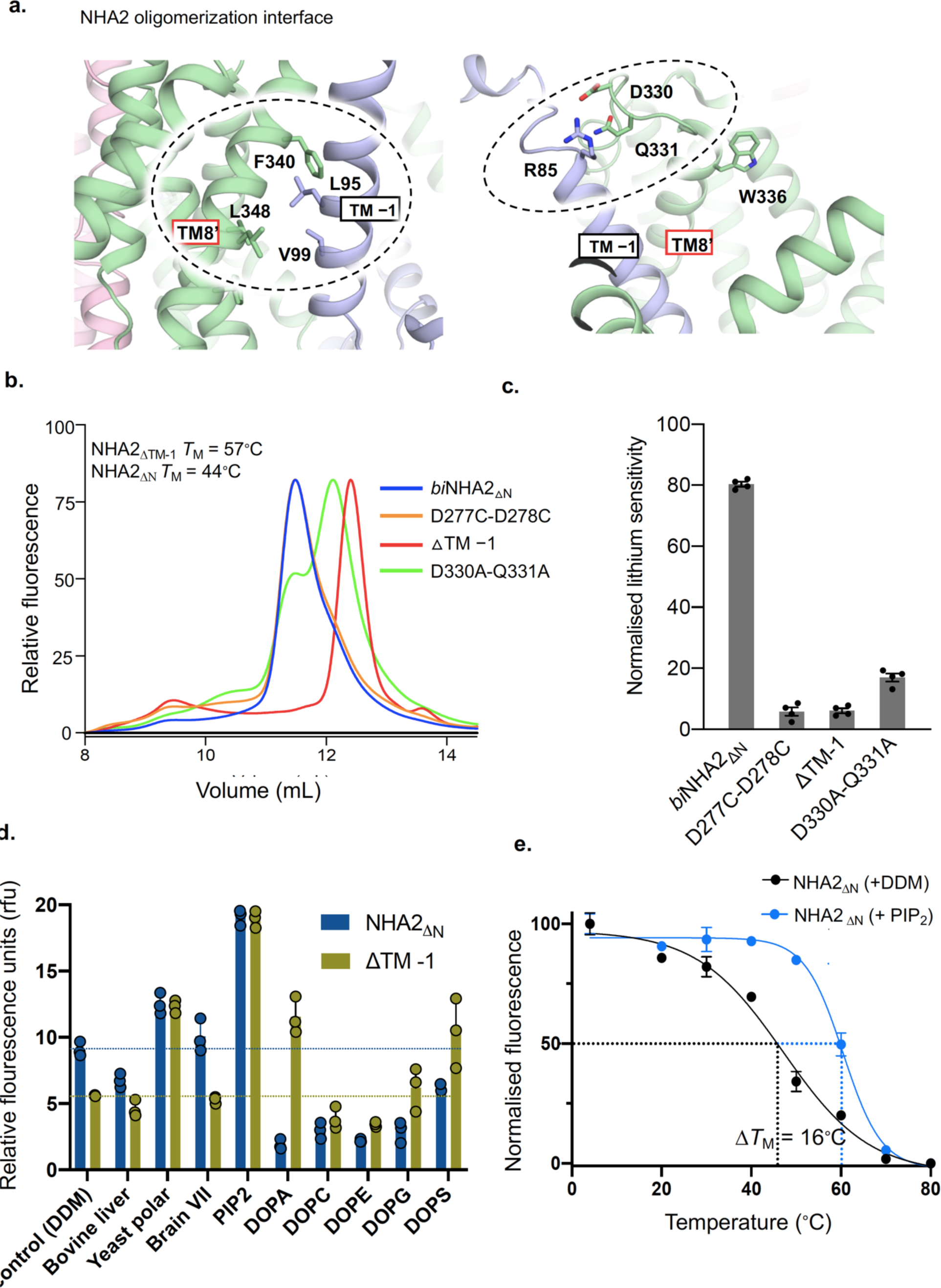
NHA2 oligomerization and lipid preferences. **a.** *left:* Cartoon representation of dimeric NHA2 from the side and coloured as in Fig. 1c. and a zoomed in focus on the oligomerization contacts formed between TM8 in one protomer (green sticks, labelled) and TM –1 on the other protomer (blue sticks, labelled). *right*: As in left panel, but showing the polar contacts between strictly-conserved D330, Q331 residues in the TM8-TM9 loop of one protomer (green sticks, labelled) and Arg85 in TM –1 of the other protomer (blue sticks, labelled). **b.** Representative FSEC traces of DDM-CHS solubilised *bison* NHA2 *Δ*N (blue) containing membranes that were isolated after heterologous expression in the salt-sensitive yeast strain *S. cerevisiae* Ab11c strain^12, 22^. B*ison* NHA2 *Δ*N mutations and the TM -1 deletion were similarity measured as labelled. **c.** Normalised sensitivity of *bison* NHA2 *Δ*N and derived constructs grown in the Ab11c strain with 20 mM LiCl (see ED Fig. 2 and Supplementary Fig. 2 for cell growth in the presence of different lithium concentrations and non-induced controls). In all experiments errors bars, s.e.m.; n = 4 independent cultures. **d.** Thermal stabilization of DDM-purified dimeric *bison* NHA2 *Δ*N-GFP (blue bars) and NHA2 *Δ*TM-1 (green bars) by lipids. Normalized mean fluorescence after heating (*T*M + 5°C) and centrifugation in presence of either the detergent DDM or DDM solubilised lipids. Data presented are mean values ± data range of n = 3 experiments (see Methods). **e**. Thermal shift stabilization of purified dimeric NHA2 *Δ*N-GFP in the presence of DDM addition (black) compared to PIP2 in DDM free (blue). Data presented are normalized mean fluorescence as mean values ± data range of n = 3 technical repeats; the apparent *T*M was calculated with a sigmoidal 4-parameter logistic regression function; the average ΔTM presented is calculated from n = 2 independent titrations.

**Fig. 4.**
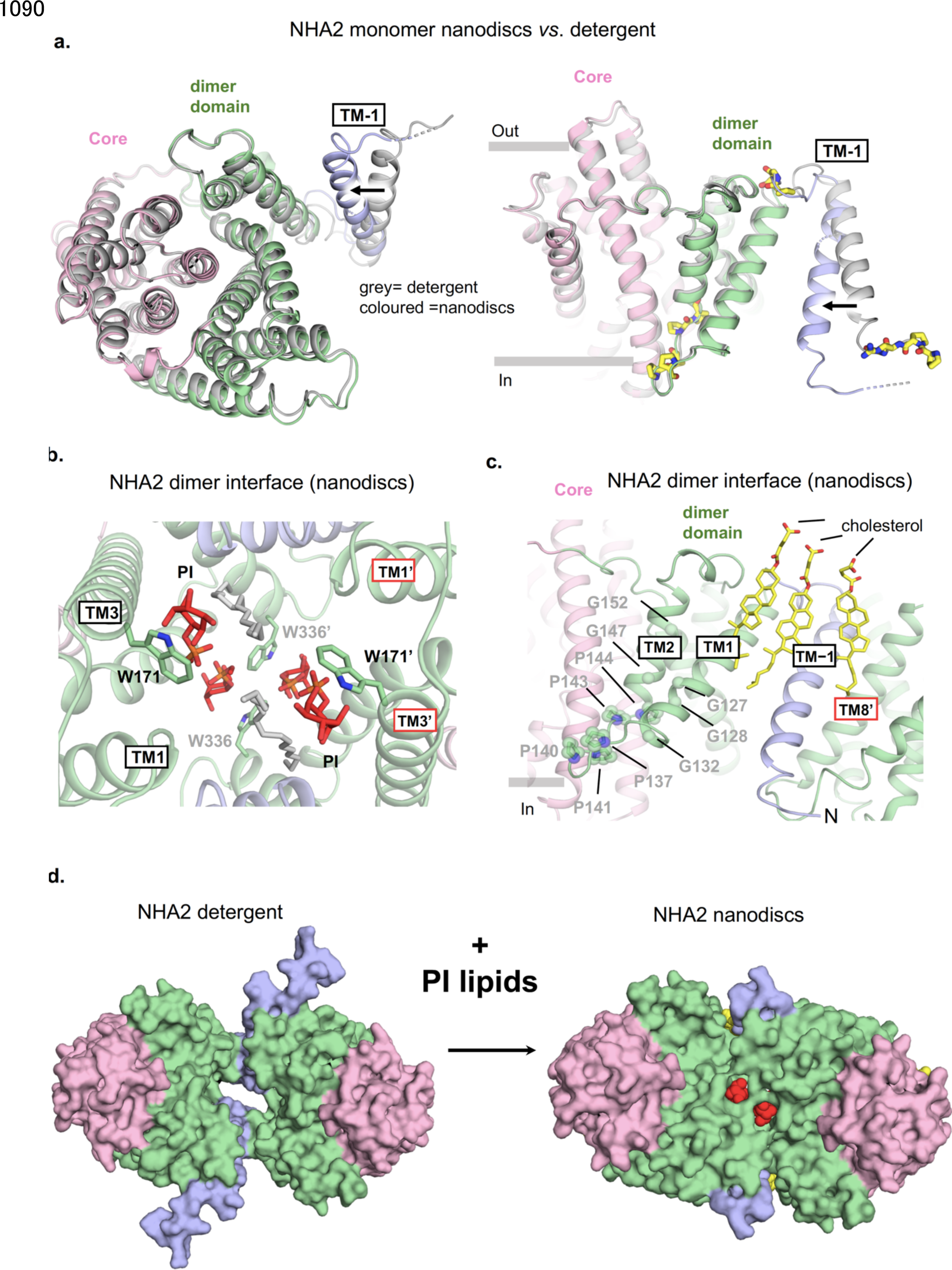
The cytosolic gap is closed by lipids in cryo EM NHA2 structure in nanodiscs. **a.** *left:* Cartoon representation of the 14-TM NHA2 monomer in nanodiscs from the intracellular side and coloured as in Fig. 1c. that has been superimposed onto the NHA2 structure in detergent in grey that highlights the movement of TM –1. *right*: As in the left panel from the side and highly-conserved including proline residues in yellow stick form (see ED Fig. 1). **b.** Cartoon representation of the NHA2 homodimer in nanodiscs from the intracellular side and zoomed in on the remodelled interface with the sugar headgroup lipids (red, orange) coordinated by tryptophan residues at the protomer interface; lipid tails that could be modelled are shown in grey. **c**. As in b. showing the side-view of the NHA2 oligomerization and highlighting cholesterol lipids (stick form, yellow) interact at the dimerization interface. Notably, the readjustment of TM –1 in nanodiscs is facilitated by highly-conserved proline and glycine residues that were shown in a. and connects to proline/leucine loop proceeding TM2 that is only 12 amino acids long. **e**. Surface representation of the *bison* NHA2 *Δ*N structure from the cytoplasmic side in detergent (left) and in nanodiscs mixed with PI lipids (right).

**Table 1.** Data collection, processing and refinement statistics of *bison* NHA2 structures.

|  | Detergent | Detergent | Nanodiscs |
| --- | --- | --- | --- |
|  | Outward facing | Outward facing | Outward facing |
|  | w/o extended helix | with extended helix | w/o extended helix |
|  | EMDB-13162 | EMDB-13161 | EMDB-13163 |
|  | PDB-7P1J | PDB-7PII | PDB-7P1K |
| Data collection and processing |  |  | 130,000 |
| Magnification | 165,000 |  | 300 |
| Voltage (kV) | 300 |  | 63.5 |
| Electron exposure (e-/Å<sup>2</sup>) | 80 |  | 0.6 – 2.2 |
| Defocus range (µm) | 0.7 - 2.5 |  | 0.68 |
| Pixel size (Å) | 0.83 |  | C1 |
| Symmetry imposed | C1 |  | 3,055,998 |
| Initial particle images (no.) |  |  | 362,665 |
| Final particle images (no.) | 1,326,947 |  |  |
|  | 269,412 |  |  |
| Map resolution (Å) | 3.04 | 3.15 | 3.58 |
| FSC threshold | 0.143 | 0.143 | 0.143 |
| Map resolution range (Å) | 2.9 – 4.6 | 3.0 – 4.8 | 3.2 - 4.6 |
| Refinement |  |  |  |
| Initial model used (PDB code) | 4cz9 |  |  |
| Model resolution (Å) | 3.8 | 4.0 | 3.1 |
| FSC threshold | 0.143 | 0.143 | 0.143 |
| Map sharpening B factor (Å<sup>2</sup>) | -80.3 | -58.2 | -100 |
| Model composition | 6502 | 6644 | 6834 |
| Non-hydrogen atoms | 868 | 884 | 866 |
| Protein residues |  |  | PI : 4 |
| Ligands |  |  | CHS : 8 |
| B factors (Å<sup>2</sup>) | 151.99 | 185.09 | 33.48 |
| Protein |  |  | 34.09 |
| Ligand |  |  |  |
| R.m.s. deviations | 0.004 | 0.003 | 0.009 |
| Bond lengths (Å) | 0.659 | 0.642 | 1.744 |
| Bond angles (°) |  |  |  |
| Validation | 2.5 | 2.34 | 2.52 |
| MolProbity score | 10.73 | 8.60 | 12.49 |
| Clashscore | 4.73 | 5.2 | 7.90 |
| Poor rotamers (%) |  |  |  |
| Ramachandran plot |  | 95.21 | 96.50 |
| Favored (%) | 93.14 | 4.79 | 3.5 |
| Allowed (%) | 6.86 | 0 | 0 |
| Disallowed (%) | 0 |  |  |

Unexpectedly, *bison* NHA2 *Δ*N is made up from 14-TM segments, rather than the 13-TMs observed in the mammalian Na^+^/H^+^ exchangers NHE1^9^ and NHE9^10^, and bacterial Na^+/^H^+^ antiporters NapA^24^, *Mj*NhaP^25^ and *Pa*NhaP^26^ or the 12-TMs observed in NhaA ^27^ (Fig. 1c-d, Extended Data Fig. 5a). To facilitate comparison to the 13-TM members the first helix in *bison* NHA2 *Δ*N was designated TM –1. The additional N-terminal helix TM –1 is domain-swapped in the dimer domain, and mediates homodimerization by interacting with TM8 and the TM8-TM9 loop from its neighbouring protomer (Fig. 1d). Due to the highly tilted angle of TM –1 and length of the connecting loop to TM1, the NHA2 protomers are held apart by ∼ 25 Å on the intracellular side to generate a homodimer with a large, predominantly hydrophobic interface of ∼5,000Å^2^ (Fig. 1c, Extended Data Fig. 6a-b). The extracellular ends of TM3 are positively-charged and proceeding TM –1 we observe cryo EM density for a helix extension that is also highly positively-charged and present on the intracellular face (Fig. 1d, Extended Data Fig. 3, Extended Data Fig. 4b, Extended Fig. 6a, b). Whilst NHA2 *Δ*N dimerizes very differently to any Na^+^/H^+^ exchanger observed to date, the NHA2 *Δ*N monomer shows high structural conservation to the outward-facing monomer of NapA (Fig. 1e and Extended Data Fig. 5b). Indeed, the NHA2 *Δ*N structure adopts a similar outward-facing conformation to NapA even if the homodimer is assembled very differently (Fig. 1f).

The characteristic feature of the “NhaA-fold” is the two discontinuous helices TM5a-b and TM12a-b in the 6-TM core ion-transport domain, which contain unwound regions that cross over each other near the centre of the membrane (Fig. 2a) ^27, 28^. Proximal to these extended helix break-points are oppositely-charged residues that neutralize the half-helical ends of TM5a and TM12b dipoles (Fig. 2a)^13^. In NHA2, a lysine residue Lys459 replaces the strictly-conserved arginine residue found in all 13-TM-members^13^, whilst Thr461 replaces the negatively-charged aspartate- or glutamate-containing residue (Fig. 2b, c). The substitution of Thr461 to alanine remained functional in *S. cerevisiae* complementation assays, whereas either Thr461 to glutamic acid or Lys459 to arginine were both non-functional under high Li^+^-salt stress, indicating that these amino acid differences have evolved to be optimal for NHA2 function (Extended Data Fig. 2c, Supplementary Fig. 1, Supplementary Fig. 3b). The absence of acidic residues in both the helix-break position and in the funnel, means that the cavity is less negatively-charged than in NapA (Fig. 2c, Extended Data Fig. 6b).

Previously, MD simulations of NapA have shown that an acidic residue in the helix-break position (Glu333) is important for initially attracting Na^+^ ions to the ion-binding aspartates, and consistently in MD simulations of NHA2 very few Na^+^ ions spontaneously reached the ion-binding aspartate Asp277 and Asp278 residues (Extended Data Fig. 7). During a total simulation time of 1.5 µs, only one spontaneous binding event and one partial binding event were observed (Supplementary Fig. 6, Extended Data Table 1) whereas in MD simulations of outward-facing NapA tens of events were recorded over a similar time period ^29^. Moreover, in simulations with either aspartate or both protonated (charge neutral), sodium ions did not diffuse into the funnel (Extended Data Fig. 7), suggesting that sodium ion binding requires both aspartates to be negatively charged to provide an electrostatically favourable environment. In contrast to the extracellular surface, the cytoplasmic surface of the core domain in NHA2 is highly negatively-charged (Extended Data Fig. 6a), which would indicate NHA2 has a preferential mode of Na^+^ uptake from the cytoplasmic side, i.e., consistent with the proposed directionality of H^+^-driven Na^+^-efflux. The extracellular half-helices TM5a and TM12b in NHA2 have also rotated and moved closer to the dimer domain than their comparable position in the outward-facing structure of NapA (Extended Data Fig. 5b). The rearrangements of these flexible half-helices are in agreement with the mobility of TM12b seen previously between different outward-facing structures of NapA, and could represent local gating differences ^29^.

The ion-binding site aspartates Asp277 and Asp278 are strictly conserved and, due to a lack of experimental map density, the side-chain rotamers were modelled based on the equivalent positions of the corresponding ion-binding aspartates from NapA (Fig. 2c, Extended Data Fig. 1). In MD simulations, a potential sodium ion binding event was captured that involved Asp277, Asp278, water molecules, and the backbone carbonyl oxygen atom of Val243 in the broken stretch of TM5a-b (Extended Data Fig. 7). In the simulations, the side chains of Asp278 and Asp277 easily adapt to the presence of a sodium ion by switching to a different rotameric state. Based on sequence conservation, mutational analysis and the initial monomeric NhaA crystal structure, both aspartates were initially thought to be the two proton carriers conveying electrogenicity^14, 27, 30, 31^. However, subsequent NhaA^32^ and NapA^24^ crystal structures in combination with functional analysis^33^ and MD simulations^34^ have proposed that a lysine residue forming a salt-bridge to one of the aspartate residues is the second proton carrier ^32, 34^. In NHA2, Asp277 is located ∼5 Å from the arginine residue Arg431, rather than the lysine residue found in electrogenic NhaA and NapA members that forms a salt-bridge to the equivalent aspartate (Fig. 2c). Due to the distance of Arg431 from Asp277 and its high pKa, it is unlikely Arg431 could act as a proton carrier in NHA2, i.e., it cannot bind and release protons at a neutral pH. Consistently, NHA2 activity is thought to be electroneutral as transport is unaffected by the collapse of the membrane potential in cells, oocytes and proteoliposomes ^16, 17, 33^.

To robustly rule out that Asp277 cannot act as a proton carrier, NHA2 was reconstituted into liposomes for solid-supported membrane (SSM)-based electrophysiology recordings, which is a more sensitive technique than patch-clamped electrophysiology for low turnover transporters^35^. SSM-based electrophysiology can also detect pre-steady state currents of the half-reaction *i.e.*, the binding and transport of Na^+^(Li^+^) across the membrane in electroneutral transporters as Na^+^ accumulates at a much faster rate than counter H^+^ efflux in the absence of *Δ*pH ^36^. Peak currents at symmetrical pH values were observed for *bison* NHA2 *Δ*N upon increasing concentrations of Na^+^ that were 20 to 50- times higher than the signal obtained from the “dead” ion-binding aspartate mutant Asp277Cys-Asp278Cys (Fig. 2d). Consistent with actual NHA2 *Δ*N functional activity, currents were pH dependent and proportional to the lipid-protein-ratio (LPR), whereas the dead ion-binding aspartate showed no differences to the signals obtained from empty liposomes (Fig. 2d and Extended Data Fig. 8a-c). Importantly, the peak currents were pre-steady state and their positive amplitudes are entirely consistent with electroneutral transport^36^, rather than electrogenic signals, which show amplitudes in the opposite direction^35, 37^. Moreover, as expected for pre-steady state currents, NHA2 activity was significantly more diminished when the pH was lowered on the outside as compared to the inside of the liposomes (Extended Data Fig. 8d), *i.e.,* H^+^ on the outside competes with Na^+^ (Li^+^) for the same ion-binding site and lowers the rate of Na^+^ (Li^+^) translocation. The *∼*30% lowering of transient Na^+^ currents when the pH is lower on the inside than outside is because the outwardly-directed *Δ*pH likely speeds up H^+^-translocation, which neutralises the signal from Na^+^ influx (Extended Data Fig. 8d). The binding affinities of *bison* NHA2 *Δ*N for Na^+^ (*K*D = 48 mM) and Li^+^ (*K*D = 6.1 mM) at pH 7.5 were determined (Extended Data Fig. 9a)^33^. Replacing NaCl with Na^+^-glucoronate showed similar peak currents and furthermore, either KCl or K^+^-gluconate addition showed no activity (Fig. 2e), ruling out that NHA2 might transport either K^+^ or Cl^-^ ions as proposed in the closely-related isoform NHA1^16^ (Supplementary Fig. 2). Given NHA2 is unable to bind Na^+^ at a lysosomal pH of 4.6, the ion affinities are consistent with an outwardly-directed lysosomal pH gradient driving the uptake of Na^+^ from the neutral pH of the cytoplasm, wherein lysosomal concentrations of 100 mM Na^+^ have been reported^38^. Taken together, the structure and SSM-based electrophysiology confirms NHA2 operates as an electroneutral transporter to drive Na^+^ efflux.

In addition to Arg431 being positioned far from Asp277, the guanidinium groups of Arg431 are also forming salt-bridge interactions with a nearby glutamate Glu214 in TM4 and a backbone carbonyl oxygen located at TM12a-b breakpoint proceeding the side-chain of Trp456 (Fig. 2b). The TM12a-b breakpoint is further stabilized by interactions with Lys459 (Fig. 2b). Bioinformatic analysis has previously shown the Glu214 residue is unique to the NHA2 members^13^ (Fig. 2c), and substitutions lacking a carboxylic group failed to rescue the salt-sensitive phenotype of the host strain^39^. Consistently, alanine mutations of either Glu214, Arg431 or Lys459 severely affected complementation under Li^+^-salt stress (Extended Data Fig. 2c, Supplementary Fig. 1, Supplementary Fig. 3b). To further investigate the importance of the NHA2-specific salt-bridge between Arg431 and Glu214, single Glu214Arg and Arg431Glu mutants were generated and, as expected, showed either no or poor complementation under Li^+^ salt-stress, respectively (Extended Data Fig. 2c, Supplementary Fig. 3). Size-exclusion profiles demonstrated that the Arg431Glu was well tolerated, but Glu214Arg was poorly folded, indicating that Lys459 might be able to compensate and maintain stability in the Arg431Glu mutant (Supplementary Fig. 1a, b). Interestingly, the swapped double mutant Glu214Arg- Arg431Glu, was able to complement Li^+^ salt stress to a similar degree as NHA2 *Δ*N, but could not complement for Na^+^ (Extended Data Fig. 1c, Supplementary Fig. 1, Supplementary Fig. 3). Strikingly, SSM-based electrophysiology confirms that this salt-bridge swapped mutant has now converted NHA2 *Δ*N into a Li^+^-specific transporter as the *K*D for the mutant remained like NHA2 *Δ*N (*K*D = 3.9 mM), whereas Na^+^ addition showed no measurable binding (Extended Data Fig. 9b, c). These results provide strong evidence that the salt-bridge pairing between Glu214 (TM4) and Arg431 (TM11) is mainly structural, but their environment can clearly fine-tune cation specificity. Lastly, the presence of the bulky amino acid tryptophan (Trp456) at the end of the TM12a is quite distinct from other Na^+^/H^+^ exchanger structure that, like NapA, have a smaller, non-aromatic side-chain in this position (Fig. 2c). The mutation of Trp456 to either phenylalanine or alanine retained some complementation for Na^+^ and Li^+^ like NHA2 *Δ*N, but not under high salt-stress (Extended Data Fig. 1c, Supplementary Fig. 1, Supplementary Fig. 3, Supplementary Fig. 4). SSM-based electrophysiology showed that the Trp456Phe mutant had in fact tighter binding affinities to NHA2 *Δ*N with a 7-fold increase for Li^+^(*K*D = 0.7 mM), yet it is unclear if this results in a slower turnover. We conclude that the size of the Trp456 residue may further shape the ion-binding site and its positioning is likely to be coordinated by interactions of both Lys459 and Arg431 in the TM12a-b breakpoint (Extended Data Fig. 5c).

Oligomerization is thought to be required for functional activity in Na^+^/H^+^ exchangers and elevator proteins in general, perhaps by anchoring the scaffold domain^40^. However, this assumption has been difficult to directly assess, as the oligomerization interfaces are typically large and therefore involve extensive protein-protein contacts^40^. In the *bison* NHA2 structure, however, the homodimer is formed only between TM –1 in one protomer and TM 8 on the other, and is predominately made up of knob-into-holes packing (Fig. 3a). Thus, NHA2 *Δ*N offers a tractable system to address this fundamental question. In NHA2, the removal of TM –1 (NHA2 *Δ*TM-1) remained well-folded with a similar expression level and melting temperature (*T*M) to NHA2 *Δ*N, but migrated as a monomer, rather than a homodimer (Fig. 3b, Supplementary Fig. 1). Based on SSM-based electrophysiology the *K*D of NHA2 *Δ*TM-1 for Na^+^ was estimated to be 2-fold lower than NHA2 *Δ*N (*K*D = 80 mM), which shows the ion-binding site remains intact (Extended Fig. 9a, Supplementary Fig. 1a). However, the NHA2 *Δ*TM-1 construct was unable to complement any Li^+^-salt stress *in vivo* and was indistinguishable from the dead ion-binding aspartate mutant growth curves (Fig. 3c, Extended Fig. 2b, Supplementary Fig. 3a). In addition to TM8, the TM8-TM9 loop contains a highly conserved DQ-motif (Asp330, Gln331) that was close enough to stabilise TM –1 *via* polar interactions (Fig. 3a), but map density was insufficient to assign side-chain positioning to confirm this. Remarkably, however, an Asp330Ala-Gln331Ala double mutant was found to be enough to shift a large fraction of NHA2 *Δ*N dimers into monomers as observed after detergent extraction of crude membranes (Fig. 3b). The Asp330Ala-Gln331Ala double mutant also failed to rescue growth of the host strain under high Li^+^ stress (Fig. 3c, Supplementary Fig. 3a). Taken together, we believe we provide compelling evidence that homodimerization is essential for NHA2 activity.

The additional TM –1 helix in NHA2 expands the structural-inverted repeat from the 5- TMs seen in NhaA to the 6-TMs in NapA and other 13-TM members, and now to 7-TMs in NHA2 (Extended Data Fig. 5a). The increase is the number of TM segments leads to structural differences in how the Na^+^/H^+^ exchangers dimerise, implying the dimerization domain has a functional role beyond that of a rigid “scaffold”. Indeed, in NhaA the dimer interface has evolved to bind the negatively-charged lipid cardiolipin, which is required for homodimerization^41–43^ and *in vivo* functional activity^44^. More recently, it was shown that the negatively-charged lipid PIP2,3 also stabilises NHE9 *via* a unique loop domain located at the dimerization interface^10^. Given the large and predominantly hydrophobic gap between protomers, we reasoned that lipids would bind to stabilise the interface and, as such, we screened a range of different lipids using a GFP-based thermal-shift assay (GFP-TS)^42^, which we had previously shown could reliable detect specific lipid-protein interactions to NhaA and NHE9 proteins ^10, 42^. Whilst the NHA2 TM-1 was found to be stabilised by all negatively-charged lipids, the NHA2 *Δ*TM-1 construct was selectively stabilized by phosphoinositide containing lipids (Fig. 3e), indicating the physiological dimer prefers this lipid type (Fig. 3f). To understand the molecular basis for lipid regulation, NHA2 *Δ*N was reconstituted with phosphoinositides (PI) lipids into nanodiscs and the cryo EM structure determined to 3.5 Å resolution (Table 1, Extended Fig. 10, Extended Data Fig. 11, Methods).

Overall, the cryo EM maps of NHA2 *Δ*N in nanodiscs are of high quality and showed similar ion-binding site interactions (Extended Data Fig. 11, Extended Data Fig. 12a). The main differences between the structure of NHA2 *Δ*N in detergent and nanodiscs, was that in nanodiscs the intracellular end of TM –1 had moved significantly inwards by some 10 Å (Fig. 4a). Interestingly, the readjustment of TM –1 completely closed the large gap between the protomers on the inside (Extended Data Fig. 12b, Supplementary Video. 1a- c). Manual placement of the NHA2 *Δ*N detergent structure into nanodiscs cryo EM maps, shows that there was plenty of space between the protein and nanodiscs (Extended Data Fig. 12c), i.e., indicating the conformational difference was not simply caused by the size of the nanodiscs. Rather, density consistent with PI lipids are now located at the dimerization interface, sandwiched between positively-charged surfaces and tryptophan residues (Fig. 4b, Extended Data Fig. 6, Extended Data Fig. 11). Tryptophan residues located at the end of TM3 and TM8 on both sides of the dimer domain are forming direct lipid-protein interactions (Fig. 4b, Extended Data Fig. 12d). We speculate that the coordination of PI lipids by the inward movement of TM8 moves the TM8-TM9 loop containing the DQ motif that catalyses the rearrangement of TM –1 (Extended Data Fig. 12e). The stabilization of TM3 on the extracellular side, by interaction of lysine and tryptophan residues to the PI headgroups, may further stabilize the outward-cavity by an interaction formed between Arg176 (TM3) and Glu406 (TM10) residues (Extended Data Fig. 12d, Extended Data Fig. 1). Indeed, the substitution of Arg176 to alanine was well-folded, but abolished Li^+^-complementation, whereas an alanine mutant of His169, located one turn away, retained growth to a similar level as *bison* NHA2 *Δ*N (Extended Data Fig. 12d, Supplementary Fig. 1, Supplementary Fig. 3c). Moreover, although cholesterol hemisuccinate (CHS) was used throughout purification of NHA2 in both detergent and nanodisc preparations, intriguingly, we find that cholesterol lipids are only clearly visible in the nanodisc NHA2 homodimer maps around TM –1 on the extracellular side (Methods, Fig. 4c, Extended Data Fig. 11). Thus, cholesterol may further aid to stabilise the compacted NHA2 homodimer.

We considered that the detergent NHA2 *Δ*N detergent structure might represent a less- physiological homodimer, but this conclusion would ignore the fact that NHA2 has evolved to have an additional N-terminal TM segment in the first place. Indeed, dynamic elastic modelling of NHA2 *Δ*N in nanodiscs shows that TM –1 can spontaneously adopt the position in the detergent structure (Extended Data Fig. 13a, Supplementary Video. 2a). Furthermore, the adjoining TM1 loop is highly mobile, and this propagates movements across the whole dimerization domain (Extended Data Fig. 13b, Supplementary Video. 2b). The requirement for TM –1 mobility is further consistent with highly-conserved proline and glycine residues located in the loop between TM-1 and TM1 and a cluster of five proline residues at the end of TM1 that proceeds TM2 (Fig. 4a, c). The modelled N-terminal extension helix is also highly positively charged (Extended Data Fig. 6a, Extended Data Fig. 12b), and is reminiscent of the positively-charged extension helix seen in the oligomeric betamine transporter BetP, which binds to negatively-charged lipids in response to a change in cell volume^45^. As such, it possible that TM –1 mobility and oligomerization might further be influenced by the extension helix, and leads to the question if NHA2 might have a role in volume sensing, as seen in other NHE proteins^2^.

In summary, the cryo EM structure of NHA2 reveals the structural similarity to the bacterial electrogenic Na^+^/H^+^ antiporters, yet the structure and SSM-based electrophysiology demonstrates how the ion-binding aspartates are coupled differently to carry out electroneutral transport and fine-tune ion affinities. We propose that Na^+^ uptake will be preferred on the cytoplasmic side, consistent with outwardly-directed H^+^ gradient driving the uptake of Na^+^ into lysosomes. Notably, Na^+^-dependent amino acid transporters of the SLC38 family^46^ are crucial for the regulation of mTOR complex 1 (mTORC1)^38, 47^ in lysosomes, and it’s likely that the Na^+^ outwardly-directed gradient is established in part by NHA2 activities. Lastly, NHA2 has evolved an additional N-terminal helix that gives rise to a homodimer that is sensitive to the lipid composition, highlighting the importance of evolving transporter oligomerization to regulate ion-driven transport. At a general level, we provide strong evidence that oligomerization is essential for optimal transport activity, and our molecular insights indicate why some secondary-active transporters may have evolved into elevator proteins.

## Materials and Methods

### NHA2 structural selection using fluorescence-based screening

NHA2 genes from *Homo sapiens (human)*, *Mus musculus*, *Bos taurus* and *Bison bison* were synthesized and cloned into the GAL1 inducible TEV-site containing GFP-TwinStrep-His8 vector pDDGFP3, and transformed into the *S. cerevisiae* strain FGY217 (MAT*α*, ura3–52, lys2Δ201, and pep4Δ) as previously described^48, 49^. The highest yielding construct were overexpressed in 2L cultures, cells harvested, and membranes isolated and solubilized with 1% (w/v) Dodecyl Maltoside (DDM, Glycon). Subsequently, the mono-dispersity of the detergent-solubilized protein product was assessed using fluorescence-detection size-exclusion chromatography (FSEC) using a Shimadzu HPLC LC-20AD/RF-20A (488 nmex, 512 nmem) instrument and Superose 6 10/300 column (GE Healthcare) in 20 mM Tris-HCl, pH 7.5, 150 mM NaCl, 0.03 % (w/v) *n*-Dodecyl β-D-maltoside (DDM). The thermostability of the highest expressing and most mono-disperse candidate constructs was determined as previously described^42, 49, 50^.

All NHA2 variants were generated with a standard PCR-based strategy and the final *bison* NHA2 ΔTM sequence (69 to 535) was as follows with TEV cleavage sequence underlined: MRNQDERAAHKDSEPSTEVNHTASSYQGRQQETGMNLRGIDGNEPTEGSNLLNNNEKMQETPAEPNHLQRRRQIHACPPRGLLARVITNVTMVILLWAVVWSVTGSECLPGGNLFGIIMLFYCAIIGGKLFGLIKLPTLPPLPPLLGMLLAGFLIRNVPVISDNIQIKHKWSSALRSIALSVILVRAGLGLDSNALKKLKGVCVRLSLGPCLIEACTSAVLAYFLMGLPWQWGFMLGFVLGAVSPAVVVPSMLLLQEGGYGVEKGIPTLLMAAGSFDDILAITGFNTCLGMAFSTGSTVFNVLKGVLEVIIGVVTGLVLGFFIQYFPSSDQDKLVWKRAFLVLGLSVLAVFSSTYFGFPGSGGLCTLVTAFLAGRGWASTKTDVEKVIAVAWDIFQPLLFGLIGAEVLITALRPETIGLCVATLGIAVLIRILVTYLMVCFAGFNIKEKIFISFAWLPKATVQAAIGSVALDTARSHGEKQLEGYGMD VLTVAFLSIIITAPVGSLLIGLLGPRLLQKAEQNKDEEDQGETSIQV<u>ENLYQFG</u>

### NHA2 purification

*S. cerevisiae (*FGY217) was transformed with the respective vector and cultivated in 12l cultures -URA media at 30°C 150 rpm in Tuner shaker flasks using Innova 44R incubators (New Brunswick). At an OD600 of 0.6 AU protein overexpression was induced by addition of galactose to final concentration of 2% (w/v) and incubation continued at 30°C, 150 rpm. Cells were harvested after 22 h by centrifugation (6,000 rpm, 4°C, 10 min), resuspended in cell resuspension buffer (CRB, 50 mM Tris-HCl pH 8.0, 1 mM EDTA, 0.6 M sorbitol) and subsequently lysed by mechanical disruption as previously described^48^. Cell debris was removed by centrifugation (4,400 rpm, 4°C, 20 min), and from the resulting supernatant membranes were subsequently isolated by ultracentrifugation (100,000 g, 4°C, 1 h) and homogenized in membrane resuspension buffer (MRB 20 mM Tris-HCl pH 7.5, 0.3 M sucrose).

For samples used for structural studies in detergent by cryo-EM, membranes were solubilized in solubilization buffer (1% (w/v) DDM (Glycon), 0.2% (w/v) cholesteryl hemisuccinate (CHS, Sigma-Aldrich), 20 mM Tris-HCl pH 8.0, 150 mM NaCl), during mild agitation for 1h at 4°C and subsequently cleared by ultracentrifugation (100,000 g 4°C, 1h). The supernatant was incubated for 2h at 4°C with 5 ml of Ni-NTA agarose (Qiagen) pre-equilibrated in wash buffer 1 WB1, 0.1% (w/v) DDM, 0.02% (w/v) CHS, 50 mM Tris-HCl pH 8.0, 300 mM NaCl and 20 mM Imidazole). This resin was transferred into a gravity flow column (Bio-Rad) and subsequently washed with 300 ml WB1. For the elution buffer EB containing 0.1% (w/v) DDM, 0.02% (w/v) CHS, 50 mM Tris-HCl pH 8.0, 300 mM NaCl and 300mM Imidazole was used. Subsequently, the eluted protein incubated over-night at 4°C in the presence of equimolar amounts of TEV- His8 protease during mild agitation. The digested protein was collected, concentrated using 100 kDa MW cut-off spin concentrators (Amicon Merck-Millipore) and subjected to size-exclusion chromatography (SEC), using a Superose 6 increase 10/300 column (GE Healthcare) and an ÄKTA system in 20 mM Tris-HCl pH 7.5, 150 mM NaCl, 0.001% (w/v) LMNG, 0.0002% (w/v) CHS.

Samples used for SSM-based electrophysiology studies were purified as described above with the following alterations: 1. The solubilization buffer used was 20 mM Tris-HCl, pH 8.0, 150 mM NaCl 1% (w/v) DDM, 0.2% (w/v) CHS and 10% Glycerol. 2. The Protein was eluted in 20 mM Tris-HCl, pH 8.0, 150 mM NaCl, 0.03% (w/v) DDM, 0.006% (w/v) CHS, and 250 mM Imidazol and TEV-digestion was performed during dialysis against Imidazole free buffer overnight. 3. Before SEC an additional reverse His-binding step was performed using a 5 ml HisTrap™ Fast Flow (Cytiva) column to remove non-digested protein and TEV-protease. 3. SEC was performed using an EnRich650 column (Bio-Rad) on Agilent LC-1220 system in 20 mM Tris–HCl pH 8, 150 mM NaCl. 0.03% (w/v) DDM, and 0.006% (w/v) CHS.

Samples used for nanodisc reconstitution and thermostabilization assays were purified as described except the protein was not subjected to TEV digestion.

### Determination of the lipid preferences of NHA2_ΔN_ and NHA2ΔTM-1 by GFP-based thermal shift (GFP-TS) assay

Characterization of thermostability and lipid thermal stabilization of NHA2*ΔN* and NHA2*Δ*TM-1 was determined as previously described^42^. In brief, GFP-fusions of NHA2*ΔN* and NHA2*Δ*TM-1 were purified as described previously in buffer containing 0.03% DDM 0.006% CHS. Samples were diluted on ice to a final concentration of ∼ 0.05 mg/ml in buffer containing 20 mM Tris-HCl pH 8, 150 mM NaCl, 1% (*w*/*v*) DDM and 1% (*w*/*v*) *β*-D-Ocytlglucoside (Anatrace). Sample aliquots of 100 μl were heated for 10 min over a temperature range of 20–80°C in a PCR thermocycler (Veriti, Applied Biosystems) and heat-denatured material pelleted at 18 000 × *g* for 30 min at 4°C. To avoid pellet contamination 80 μl of the supernatant were transferred to a 96-well plate (Greiner) and fluorescence was measured using a Fluoroskan plate reader (ThermoFisher, Skanlt software 6.0.2). The apparent *T*m was calculated by plotting the average GFP fluorescence intensity of three technical repeats per temperature and fitting the curves to a sigmoidal logistic regression in GraphPad Prism software. To screen for differential lipid stabilization, the GFP-TS protocol was utilized with the modification that the protein was incubated for 30 min at 4°C with the individual lipids at a final concentration of 3 mg/ml (1 mg/ml for PIP2) in 1% (w/v) DDM and these samples were heated at a single temperature (Tm of the respective construct + 5°C). Stock solutions of the respective lipids DOPA, DOPC, DOPE, DOPG, Liver Total Lipid Extract (Bovine), Yeast Polar Lipid Extract (*S. cerevisiae*) (Avanti), Brain Extract from bovine brain Type VII (Sigma-Aldirch) and PIP2, (Larodan) were prepared by solubilization in 10% (*w*/*v*) DDM to a final concentration of 30 mg/ml (10 mg/ml for PIP2) overnight at 4°C with mild agitation.

### Preparation of *bison* NHA2_ΔN_ nanodisc reconstituted sample

MSP1E2 with an N-terminal hexa-His-tag and a TEV protease recognition site was purified as previously described^51^ with the addition of His-tag cleavage during dialysis. After cleavage and prior to storage an additional reverse His-trap step was introduced to remove His-tagged TEV protease from the sample. Yeast PI solubilized in chloroform (Larodan) was dried using a rotary evaporator (Hei-Vap Core, Heidolph Instruments GmbH) to completely remove residual chloroform. Dry lipids were thoroughly resuspended in 20 mM Tris pH 7.5, 150 mM NaCl buffer at a final concentration of 20 mM. For the nanodisc reconstitution, SEC purified *bi*NHA2-GFP fusion in buffer containing 0.03% DDM 0.006% CHS, MSP1E2 and the lipid mixture were mixed at a molar ratio of 1:5:50, respectively. The mixture was kept on ice for 30 min. The solution was then transferred to a tube containing ∼30- 50 mg of Bio-Beads (Bio-Beads SM-2, BioRad) that were previously equilibrated in 20 mM Tris-HCl pH 7, 150 mM NaCl buffer. The tube was kept on a rotary shaker at 4°C overnight. Bio-Beads were sedimented by centrifugation (1 min, 3000 × g, 4°C) and the supernatant was separated by pipetting. To remove the empty nanodiscs the mixture was then incubated with equilibrated Ni-NTA (∼1 ml per mg of initial protein concentration used) in batch for 2-3 h on a rotary shaker. Non-bound supernatant was removed by subsequently washing of the resin with 5 CV of 20 mM Tris pH 7.5, 150 mM NaCl buffer and 1 CV 20 mM Tris pH 7.5, 150 mM NaCl, 25 mM Imidazol buffer. Bound nanodisc incorporated protein was eluted in 2 CV 20 mM Tris pH 7.5, 150 mM NaCl, 250 mM Imidazol buffer. Protein amount was determined by GFP fluorescence on Platereader. TEV protease was added 2:1 (w/w) and the elute was dialysed against 20 mM Tris pH 7.5, 150 mM NaCl buffer overnight. The mixture was then concentrated to an appropriate volume using a centrifugal filter (Amicon, Merck-Millipore) with a cut-off of 100 kDa and subjected to SEC using an EnRich 650 column (Bio-Rad) and an Agilent LC-1220 system in 20 mM Tris–HCl pH 7.5, 150 mM NaCl. The peak fraction corresponding to the reconstituted *bi*NHA2 in nanodisc (bands identified by SDS PAGE) was concentrated to 2.7 mg/ml for later use in Cryo-EM experiments.

### Complementation assay for mammalian NHA2 against salt-stress in yeast

*S. cerevisiae* strain Ab11c (*ena1-4Δnhx1Δnha1Δ*) lacking endogenous Na^+^(Li^+^) efflux proteins^22^ was used as an host for the expression of *bison* and *human* NHA2 and constructs. To confirm expression and folding of NHA2 constructs in the Ab11c strain, heterologous expression condition were repeated as previously described for the FGY217 strain and membranes isolated for FSEC analysis^48^. In brief, cells were cultivated in 10 ml of culture in 50 ml aerated capped tubes -URA media at 30°C 150 rpm. At an OD600 of 0.6 AU protein overexpression was induced by addition of galactose to final concentration of 2% (w/v) and incubation continued at 30°C, 150 rpm. The cells were harvested after 22 h by centrifugation (3,000 rpm, 4°C, 5 min), resuspended in cell resuspension buffer (YSB, 50 mM Tris–HCl (pH 7.6), 5 mM EDTA, 10% glycerol, 1× complete protease inhibitor cocktail tablets). To estimates the NHA2 expression levels from whole cells, the cell suspension was transferred to a 96-well plate optical bottom plate and measured GFP fluorescence emission at 512 nm by excitation at 488 nm in a Fluoroskan plate reader as previously described^48^. The cells subsequently lysed by glass beads (Sigma, St. Louis, MO). Cell debris was removed and from the resulting supernatant membranes were isolated by ultracentrifugation (120,000 g, 4°C, 1 h) and resuspended YSB. The membranes solubilized with 1% (w/v) DDM. Subsequently, the mono-dispersity of the detergent-solubilized protein product was assessed using FSEC using a Shimadzu HPLC LC-20AD/RF-20A (488 nmex, 512 nmem) instrument in 20 mM Tris-HCl, pH 7.5, 150 mM NaCl, 0.03 % (w/v) DDM.

Salt-sensitive growth was monitored by inoculating 0.2 ml of -URA medium and 2% galactose final in a 96-well plate with a saturated seed culture that had been grown o/n in 2% glucose to a starting OD600 of either 0.2 for cultures with Li^+^ addition or 0.02 for Na^+^ addition. After incubation at 30°C for 48–72 h the cultures were resuspended and the OD600 measured using Spectramax plate reader (Molecular Devices).

### SSM-based electrophysiology of *bison* NHA2_ΔN_

SSM-based electrophysiology measurements were performed on SURFE²R N1 (Nanion Technologies) with liposome reconstituted protein^35^. For liposome reconstitution yeast polar lipids (Avanti) solubilized in chloroform were dried using a rotary evaporator (Hei-Vap Core, Heidolph Instruments GmbH) to completely remove residual chloroform. Dry lipids were thoroughly resuspended in 20 mM Tris pH 8, 20 mM NaCl, 20 mM KCl, 5 mM MgCl2 buffer at a final concentration of 10 mg/ml. The lipids were flash-frozen in liquid nitrogen and then thawed, in a total of eight freeze–thaw cycles. Unilamellar vesicles were prepared by extruding the resuspended lipids using an extruder (Avestin, Inc.) with 400 nm polycarbonate filters (Whatman). Vesicles were destabilized by addition of a Na-Cholate (0.65% final concentration). SEC purified protein was added to the destabilized liposomes at a lipid-to-protein ratio (LPR) of 5:1 (or otherwise stated) and incubated for 5 min at room temperature. Detergent was removed using a PD-10 desalting column, and the sample was collected in a final volume of 2.3 mL. Proteoliposomes were centrifuged at 150,000 × *g* for 30 min and resuspended to a final lipid concentration of 1 mg/ml in 20 mM Tris pH 8, 20 mM NaCl, 20 mM KCl, 5 mM MgCl2 buffer and stored at -80°C. Proteoliposomes were diluted 1:1 (v/v) with non-activating buffer (20 mM Tris at desired pH, 300 mM KCl, 5 mM MgCl2) and sonicated using a bath sonicator prior to sensor preparation ^35^.

Sensor preparation for SSM-based electrophysiology on the SURFE^2^R N1 was performed as described previously^35^ During the experiments, NHA2 was activated by solution exchange from non-activating buffer to an activating buffer containing the substrate. Usually x mM KCl is replaced by x mM NaCl or LiCl in the activating buffer. In some cases, 150 mM KCl was replaced by 150 mM choline chloride in the non-activating solution. In pH jump experiments, the activating buffer was titrated to the desired pH.

For measurements in presence of pH gradients, a double solution exchange workflow was applied: Proteoliposomes were diluted and sonicated in non-activating buffer to equilibrate the intraliposomal pH (pHi) prior to sensor preparation. During the experiment, a non-activating solution with a different pH is applied to adjust the external pH (pHo). 1 s after establishment of the pH gradient, non-activating solution was exchanged by activating solution containing 150 mM NaCl to activate NHA2 in presence of pH gradient. After the experiment, the sensor was rinsed with non-activating solution at pHi and incubated for 3 minutes to re-establish the intraliposomal pH.

### Cryo-EM sample preparation and data acquisition

#### NHA2ΔN structure in detergent

The purified *bison* NHA2*Δ*N protein was individually applied to freshly glow-discharged Quantifoil R2/1 Cu300 mesh grids (Electron Microscopy Sciences). Grids were blotted for 3.0 s or 3.5 s under 100% humidity and plunge frozen in liquid ethane using a Vitrobot Mark IV (Thermo Fisher Scientific). Cryo-EM datasets were collected on a Titan Krios G2 electron microscope operated at 300 kV equipped with a GIF (Gatan) and a K2 summit direct electron detector (Gatan) in counting mode. The movie stacks were collected at 165,000× corresponding to a pixel size of 0.83 Å at a dose rate of 7.0–8.0 e^-^ per physical pixel per second. The total exposure time for each movie was 10 s, thus leading to a total accumulated dose of 80 e^−^ Å^−2^, which was fractionated into 50 frames. All movies were recorded with a defocus range of −0.7 to −2.5 µm. The statistics of cryo-EM data acquisition are summarized in Extended Data Table 1.

#### NHA2ΔN structure in nanodiscs

The nanodisc incorporated protein was individually applied to freshly glow-discharged Quantifoil R1.2/1.3 Cu300 mesh grids (Electron Microscopy Sciences). Grids were blotted for 4 s under 100% humidity and plunge frozen in liquid ethane using a Vitrobot Mark IV (Thermo Fisher Scientific). Cryo-EM datasets were collected on a Titan Krios G2 electron microscope operated at 300 kV equipped with a GIF (Gatan) and a K3 summit direct electron detector (Gatan) in super-resolution counting mode. The movie stacks were collected at 130,000× corresponding to a pixel size of 0.68 Å at a dose rate of 14.3 e^-^ per physical pixel per second. The total exposure time for each movie was 2 s, thus leading to a total accumulated dose of 63.5 e^−^ Å^−2^, which was fractionated into 40 frames. All movies were recorded with a defocus range of −0.6 to −2.2 µm. The statistics of cryo-EM data acquisition are summarized in Extended Data Table 1.

### Image processing and model building

#### NHA2ΔN structure in detergent

Dose-fractionated movies were corrected by using MotionCorr2^52^. The dose-weighted micrographs were used for contrast transfer function estimation by CTFFIND-4.1.13^53^. The dose-weighted images were used for auto-picking, classification and reconstruction. For NHA2 datasets, approximately 1,000 particles were manually picked, followed by one round of 2D classification to generate templates for a subsequent round of auto-picking in RELION-3.0^54^. The auto-picked particles were subjected to multiple rounds of 2D classification in RELION-3.0 to remove “junk particles”. Particles in good 2D classes were extracted for initial model generation in RELION-3.0^54^.

The initial model was low-pass filtered to 20 Å to serve as a starting reference for a further round of 3D auto-refinement in RELION 3.0 or 3.1 using all particles in good 3D classes. Good 3D classes were selected and iteratively refined to yield high-resolution maps in RELION 3.0 or 3.1 with no symmetry applied. To improve the map quality, per particle CTF refinement and Bayesian polishing^55^ was performed using carried by RELION 3.0 or 3.1. and the resolution reached to 3.56 Å. The polished particle from RELION was performed Non-uniform refinement and local refinement in cryoSPARC v2.14.2^56^. After the refinement, to remove the micelle signal, signal subtraction was performed. To convert the format from RELION to cryoSPARC v2.14.2, *UCSF* pyem^57^ was used. The subtracted particle data from cryoSPARC set was performed refinement in RELION with SIDESPRITTE^58^. The overall resolution was estimated based on the gold-standard Fourier shell correlation (FSC) cut-off at 0.143. The local resolution was calculated from the two half-maps using RELION 3.1. The particles were then subjected to one round of 3D classification without alignment using a regularization parameter of 100, yielding five 3D classes. Of the five 3D classes, four classes closely resembled the outward facing structure and one class consisting of 18.3% was the ion-binding structure. The final resolution for the outward structure from 4 of 5 classes and 1 of 5 classes from ion-binding structure were 3.0 Å and 3.5 Å based on gold-standard FSC, respectively.

Homology modelling was initially performed by SWISS-MODEL using the crystal structure of NapA as a template (PDB id: 4bwz). The model was fitted as a rigid-body into the map density using MOLREP in CCP-EM suite^59^. After molecular replacement, manual model building was performed using COOT^60^. The final model underwent simulated annealing and NCS restraints using real space refinement in PHENIX^61^. The refinement statistics are summarized Extended Data Table 1.

#### NHA2ΔN structure in nanodiscs

Dose-fractionated movies were corrected using Patch motion and the dose-weighted micrographs were used for contrast transfer function estimation by Patch CTF. The dose-weighted images were used for auto-picking (Blob picker), classification and reconstruction. Initially auto-picked particles were used for one round of 2D classification to generate templates for a subsequent round of auto-picking. The auto-picked particles were subjected to multiple rounds of 2D classification, ab-initio reconstruction and hetero refinement to remove “junk particles”. All the steps mentioned above were performed in cryoSPARC v3.2.0^56^. Additionally, particle removal was performed using 2D classification without alignment in RELION 3.1^54^. To convert the format from cryoSPARC to RELION, *UCSF* pyem^57^ was used. The particle stack was reimported to cryoSPARC and subsequential heterologous refinement steps performed. The final particle stack of the best class from heterologous refinement was further processed with non-uniform refinement and multiple local refinement steps in cryoSPARC v3.2.0. The last refinement step was performed in RELION 3.1 with SIDESPLITTER^58^. The overall resolution was estimated based on the gold-standard Fourier shell correlation (FSC) cut-off at 0.143. The local resolution was calculated from the two half-maps using RELION 3.1.

### Molecular dynamics simulations

#### All-atom, explicit solvent MD simulations

NHA2 was simulated with all-atom, explicit solvent molecular dynamic (MD) simulations in a realistic model of the lysosomal membrane. Simulations were based on the NHA2*Δ*N nanodisc structure. The protein was embedded in a model for the lysosomal outer membrane, approximated by a composition of 65% PC, 20% PE, 6.5% PI, 4% SM, 4% PS, 0.5% PIP2 ^62, 63^. No detailed data on the acyl chains composition of lipids in the lysosomal membrane was available. Therefore, experimental data for red blood cell plasma membranes in ^62^ was used as a starting point: PC 16:0/18:1, PE 16:0/18:1, SM 34:1, SM 40:1, and SM 40:2 (three SM species in equal proportions 1:1:1), PS 18:2/18:0, PI 18:0/20:4, and PIP2 with CHARMM36 lipid parameters ^64^ POPC, POPE, SAPI, PSM/LSM/NSM, SLPS, and SAPI25 with a total of 203 lipids in the cytosolic leaflet and 199 in the lysosolic leaflet. The protein/membrane system was simulated with a free NaCl concentration of 150 mM. The system was built with CHARMM-GUI v1.7 ^65–67^ using the CHARMM27 force field with cmap for proteins ^68, 69^, CHARMM36 for lipids ^64^, and the CHARMM TIP3P water model. Simulation systems contained around 179,000 atoms in an hexagonal unit cell with unit cell lengths *a*=140 Å, *c*=110 Å. All titratable residues were simulated in their default protonation states at pH 7.5 as predicted by PROPKA 3.1 ^70^, with the following exceptions: The pKa of Lys459 was predicted as downshifted to ∼7 due to the interaction with Arg431 and hence it was modelled as deprotonated (neutral). Additionally, the putative binding site residues Asp277 and Asp278 were simulated in all combinations of protonation states as described below. Equilibrium MD simulations were performed with GROMACS 2021.1^71^ on GPUs. Initial relaxation and equilibration followed the CHARMM-GUI protocol ^65^ with an initial energy minimization and 6 stages of equilibration with position restraints (harmonic force constant on protein and lipids). We ran all simulations in the NPT ensemble at constant temperature (T = 303.15 K) and pressure (P = 1 bar). The stochastic velocity rescaling thermostat ^72^ was used with a time constant of 1 ps, and three separate temperature-coupling groups for protein, lipids and solvent. Semi-isotropic pressure coupling was achieved with a Parrinello-Rahman barostat ^73^ with time constant 5 ps and compressibility 4.5 × 10^−5^ bar^−1^. The Verlet neighbor list was updated as determined by gmx mdrun for optimum performance during run time within a Verlet buffer tolerance of 0.005 kJ/mol/ps. All simulations employed periodic boundary conditions and therefore Coulomb interactions were calculated with the fast-smooth Particle-Mesh Ewald (SPME) method ^74^ with an initial real-space cutoff of 1.2 nm and interactions beyond the cutoff were calculated in reciprocal space with a fast Fourier transform on a grid with 0.12-nm spacing and fourth-order spline interpolation. The balance between real space and reciprocal space calculation was optimized by the Gromacs GPU code at run time. The Lennard-Jones forces were switched smoothly to zero between 1.0 nm and 1.2 nm and the potential was shifted over the whole range and decreased to zero at the cut-off. Bonds to hydrogen atoms were converted to rigid holonomic constraints with the P-LINCS algorithm ^75^ or SETTLE ^76^ (for water molecules). The classical equations of motions were integrated with the leapfrog algorithm with a time step of 2 fs.

### Simulations

All-atom, explicit solvent simulations were performed with different protonation states of Asp277 and Asp278. We explicitly modelled all four combinations of protonation states for these two important residues. We hypothesized that the state in which NHA2 is likely to bind a sodium ion, has both Asp277 and Asp278 deprotonated and therefore negatively charged (*state 0*). As controls, we also modelled states less likely to support ion binding. In *state 1*, Asp277 is protonated (and neutral) while Asp278 remains deprotonated and negatively charged. In *state 2*, Asp277 remains deprotonated while Asp278 is protonated. Finally, the state least likely to support cation binding is *state* 3 with both aspartate residues deprotonated. Because NHA2 is a homodimer, we sampled two different charge states in a single simulation by preparing protomer A in state 0 (or 2) and protomer B in state 1 (or 3). Three independent repeats were performed by varying the initial velocities for simulations including state 0 whereas two repeats were carried out for the state 2/3 simulations (see Extended Data Table 1).

### Trajectory analysis

Analysis was carried out with Python scripts based on MDAnalysis ^77^ (distances, RMSD, RMSF). Sodium density maps were calculated with MDAnalysis from trajectories that were structurally superimposed on all C*α* atoms of protomer A (for simulation f-01-0) or all protein C*α* atoms (simulation f-23-1). Lipid density maps were calculated for all lipids, PI, and POPE in simulation f-01-0. Trajectories were aligned to all C*α* atoms atoms protein before performing density calculations. Membrane thickness was analyzed with FATSLiM ^78^. Time series of bound Na^+^ distances to carboxyl oxygen atoms in D277 and D278 were calculated for all Na^+^ ions as the shortest distance between Na^+^ and either Asp OD1, OD2 atoms. Binding of Na^+^ ions were assessed with a simple distance criterion: any Na^+^ ion within 3 Å of any carboxyl oxygen atom of Asp277 was considered bound. Molecular images were prepared in VMD 1.9.4 ^79^ and UCSF Chimera 80.

### Elastic Network Modelling (ENM) and nanodisc transition

Elastic Network Models (ENM) represent proteins as a simple network of residues (C-alphas) connected by elastic springs, so that diagonalization of the connectivity matrix renders 3N-6 eigenvectors or Normal Modes (NMs) that describe the large-scale conformation changes seen in experimental structures. To obtain an approximation to the intrinsic dynamics of NHA2, NMs were computed using the MD-refined potential ED-ENM as previously described ^10, 81, 82^ . Mid-frequency modes computed from NHA2 outward conformation were found to display elevator-like movements between the dimer and transport domains capable to partially drive the transition towards inward-like states resembling NapA inward conformation (Extended Data Fig. 13). Transition pathways between NHA2 conformations in detergent and nanodiscs were generated with coarse-grained eBDIMS Langevin simulations ^81, 82^.

## Supporting information

Extension and Supplementary Data

## Acknowledgement

We are grateful for the salt-sensitive yeast Abc11 strain from Olga Zimmermannova and to Cryo-EM Swedish National Facility at SciLife Stockholm for cryo-EM data collection as well as the Umeå Core Facility for Electron Microscopy, UCEM and the European Synchrotron Radiation Facility (ESRF). This work was funded by grants from The Swedish Research council (VR) and a European Research Council (ERC) Consolidator Grant EXCHANGE (Grant no. ERC-CoG-820187) to D.D. Research reported in this publication was supported by the National Institute of General Medical Sciences of the National Institutes of Health under Award Number R01GM118772 to O.B. MD simulations were performed using SDSC Expanse at the San Diego Supercomputing Center (allocation TG-MCB130177 to O.B.), a resource of the Extreme Science and Engineering Discovery Environment (XSEDE), which is supported by National Science Foundation grant number ACI-1548562. The authors also acknowledge Research Computing at Arizona State University for providing HPC and storage resources that have contributed to the research results reported within this paper. This work was partly supported by grants from Basis for Supporting Innovative Drug Discovery and Life Science Research (BINDS) from the Japan Agency of Medical Research and Development (AMED) (Grant no. JP20am0101079). This research was also supported by Cancerfonden Junior Investigator Award (21 0305 JIA 01 H) and Jeanssons Foundations Grants to L.O. eBDIMS simulations were performed using the Swedish National Infrastructure for Computing (allocation SNIC 2021/5-87 to L.O.).

## Data availability

The atomic coordinates and cryo-EM maps for NHA2 are available at the Protein Data Bank (PDB)/Electron Microscopy Data Bank (EMDB) databases. The accession numbers are NHA2 detergent with extension helix 7P1I/EMD-13161; NHA2 detergent without extension helix 7P1J/EMD-13162 and NHA2 in nanodiscs 7P1K/EMD-13163. The data sets generated in the current study are available from the corresponding author on reasonable request.

## Author contribution

D.D. designed the project. Cloning, expression screening and sample preparation for cryo-EM was carried out by R.M and R.F. FAB fragments used for initial crystallization trials were made by N.N and S.I. Cryo-EM data collection and map reconstruction was carried out by R.M. and R.F. Model building was carried out by R.M., R.F. and D.D. Experiments for functional analysis were carried out by R.F, S. J, A.B, and Y.C. L.O. performed ENM analysis and eBDIMS Langevin simulations. MD simulations were carried out by L.O. C.Z, and OB. All authors discussed the results and commented on the manuscript. Correspondence and request for materials should be addressed to D.D.

