## Supplementary material for "Structure and lipid-mediated remodelling mechanism of the Na^+^/H^+^ exchanger NHA2": Extension and Supplementary Data

**Extended Data Table 1: MD simulations**.

| simulation name | state-A | state-B | atoms | run length (ns) | binding -A | binding-B |
|---|---|---|---|---|---|---|
| f-01-0 | 0 | 1 | 178522 | 716.6 | strong(1) | — |
| f-01-1 | 0 | 1 | 178522 | 390.1 | — | — |
| f-01-2 | 0 | 1 | 178522 | 386.9 | partial(2) | — |
| f-23-1 | 2 | 3 | 179068 | 374.1 | — | — |
| f-23-2 | 2 | 3 | 179068 | 382.1 | — | — |

Protomers A and B were prepared with different protonation states: **state 0** (both Asp277 and Asp278 deprotonated), **state 1** (Asp277 protonated, Asp278 deprotonated), **state 2** (Asp277 deprotonated, Asp278 protonated), and **state 3** (both Asp277 and Asp278 protonated).

(1) Spontaneous binding from 349.9 ns onwards until the end of the simulation.

(2) Spontaneous partial binding from 327.0 ns onwards until the end of the simulation.

### Extended Data Fig. 1

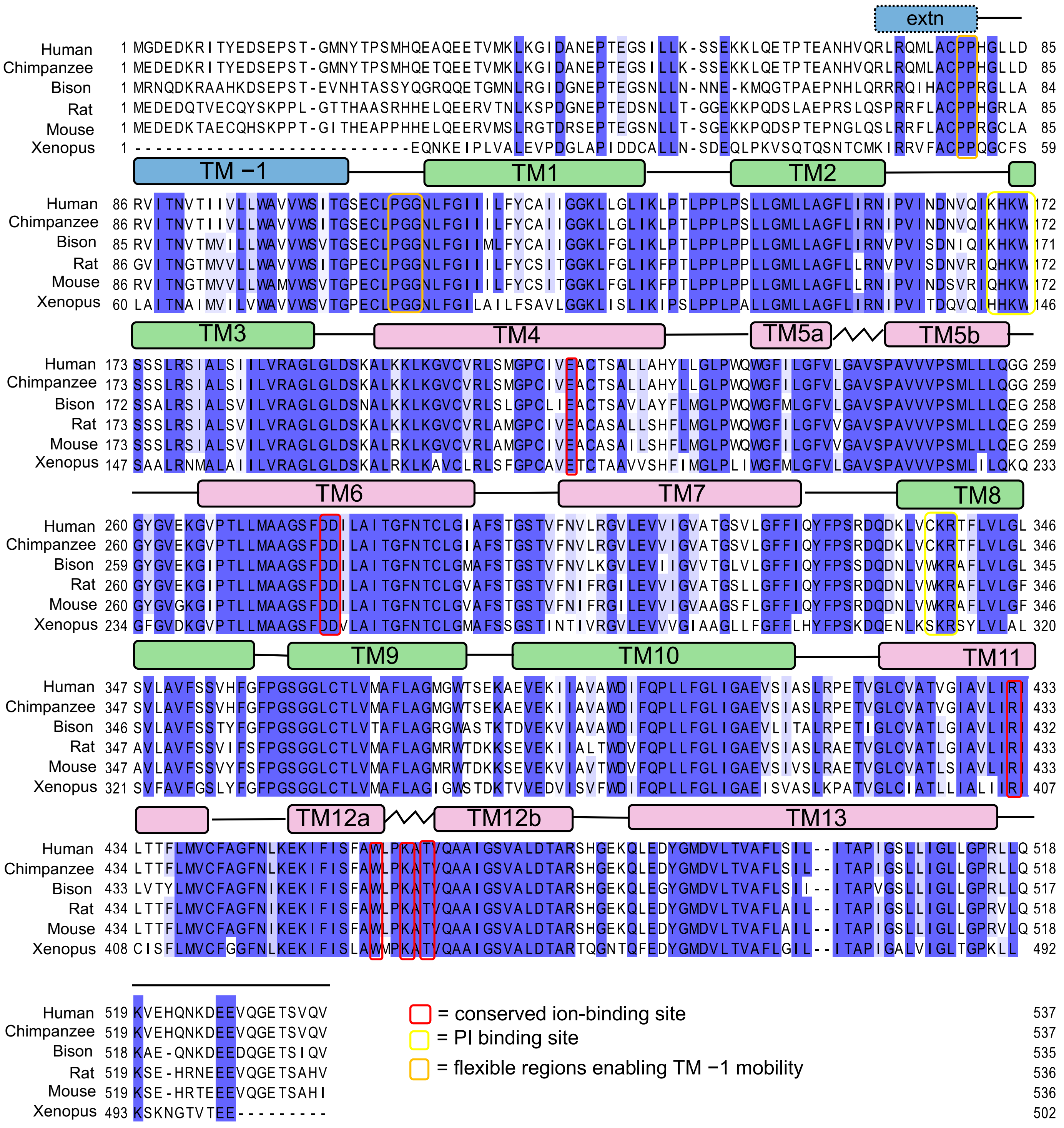

**Extended Data Fig. 1. Multiple sequence alignment of *bison* NHA2 and eukaryotic homologues.** NHA2 sequences were aligned from selected sequences of mammalian vertebrates from *Xenopus laevis* (African clawed frog) to Human (SLC9B2). Residues with over 70% sequence identity are indicated by purple background. Conserved ion-binding site residues are highlighted with a red border. Positions which have been identified to regions for lipid binding are highlighted with yellow border and proline and glycine clusters associated with TM −1 mobility with orange border. Breakpoints (s-shaped line), core 6-TM transport domain TMs (pink), dimerization domain (green) and domain-swapped helix TM −1 (blue) are indicated.

**Extended Data Fig. 2**

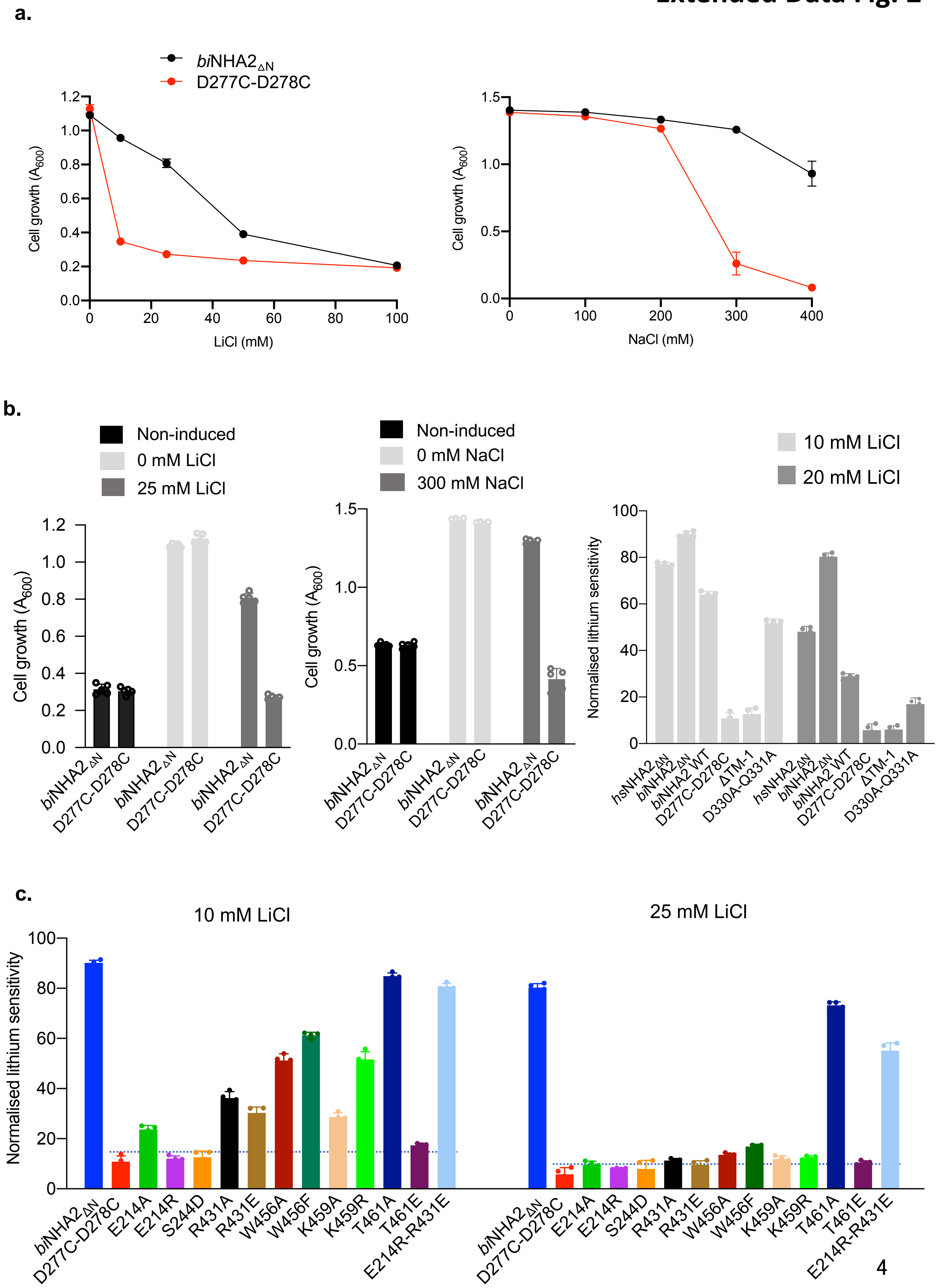

**Extended Data Fig. 2. Functional complementation of salt-sensitive yeast strain AB11c by heterologous expression of *bison* NHA2 and mutants**. **a**. Ab11c strain was transformed with *bison* NHA2$_{\Delta N}$ or the corresponding construct with both Asp277 and Asp278 have been substituted with cysteine (D277C-D278C); *human* NHA2 numbering is D278, D279. Yeast were grown as outlined in Methods in -URA media supplemented with 2% galactose and either LiCl (left) or NaCl (right) and growth was determined by optical density of the culture at 600 nm ($OD_{600}$) after 48 or 72 h at 30°C, respectively. **b**. *left:* as in a., for NHA2 constructs grown in in the absence or presence of either 0, 20 or 25 mM LiCl or 300 mM NaCl and *right*: normalised lithium sensitivity based on growth of AB11c yeast cells grown in the absence of salt-stress **c**. normalised lithium sensitivity at 10 and 25 mM LiCl concentrations as described in b. (see Supplementary Fig. 1 for expression and FSEC curves and Supplementary Fig. 3 for other $Li^+$ concentrations and non-normalized data). In all experiments described in a – c the errors bars, s.e.m.; n = 4 independent cultures.

#### Extended Data Fig. 3

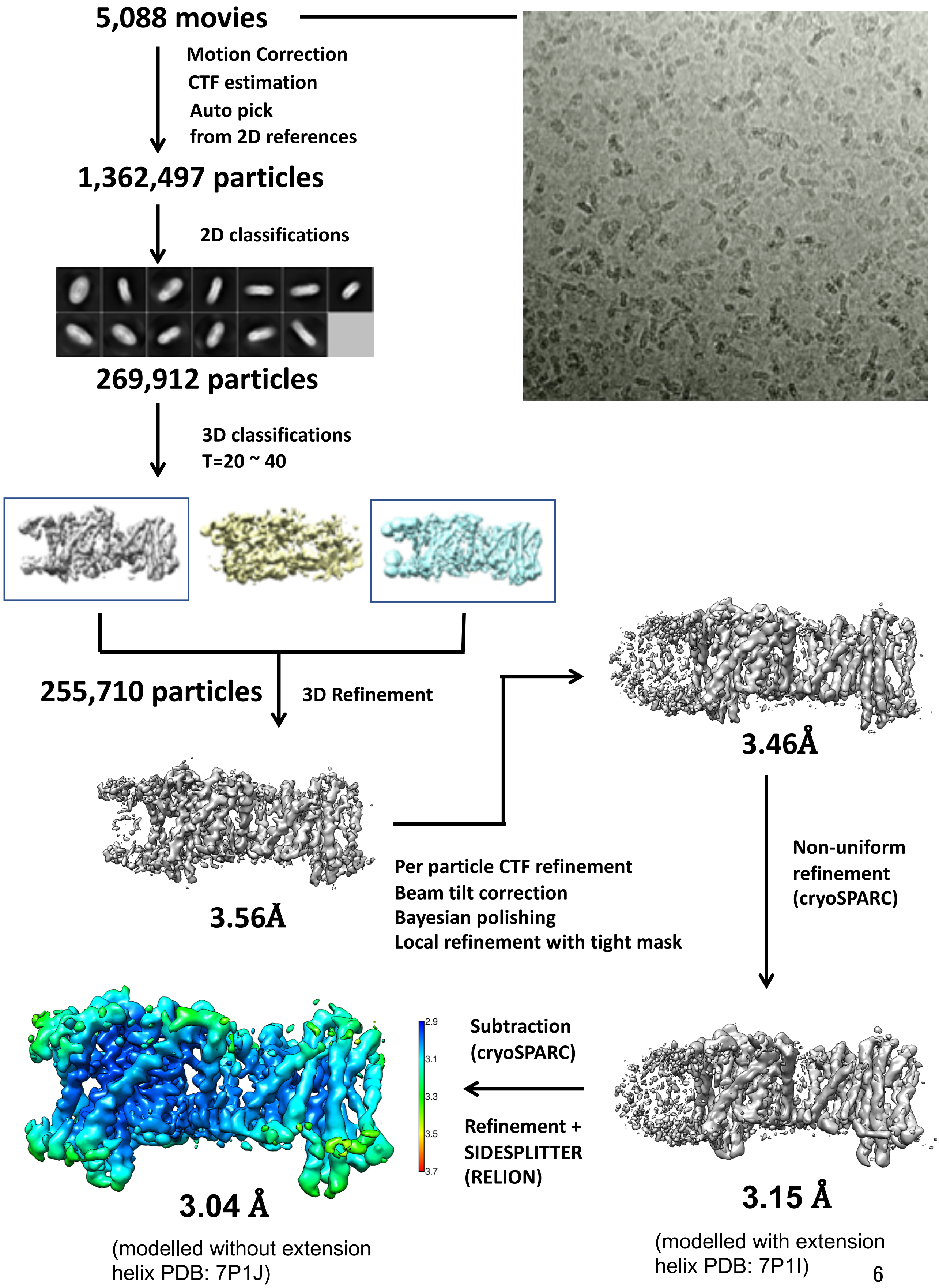

**Extended Data Fig. 3. The data-processing workflow of *bison* NHA2 $_{\Delta N}$ in detergent.** The dataset contained 5,088 movies that were corrected by MotionCor2 and CTFFind. After reference based autopicking, 1,362,497 particles were picked. Several rounds of 2D classification were performed, yielding 269,912 particles, which were subjected to 3D classification. One of the three 3D classes was selected, and it contained 255,710 particles. After several rounds of refinement with global and local search using CTF refine, polishing and masking 3.46Å was achieved. After non-uniform refinement in cryoSPARC, a resolution of 3.15Å was achieved at gold standard FSC (0.143), which was later used for modelling the N-terminal helix (after the final model was built from the next step of refinement), The final maps were obtained particle subtraction of micelle region in cryoSPARC and SIDESPLITTER in Relion a resolution of 3.04Å was achieved at gold standard FSC (0.143), with a local resolution range of 2.9 to 3.7 Å.

12 **a.**

**Extended Data Fig. 4**

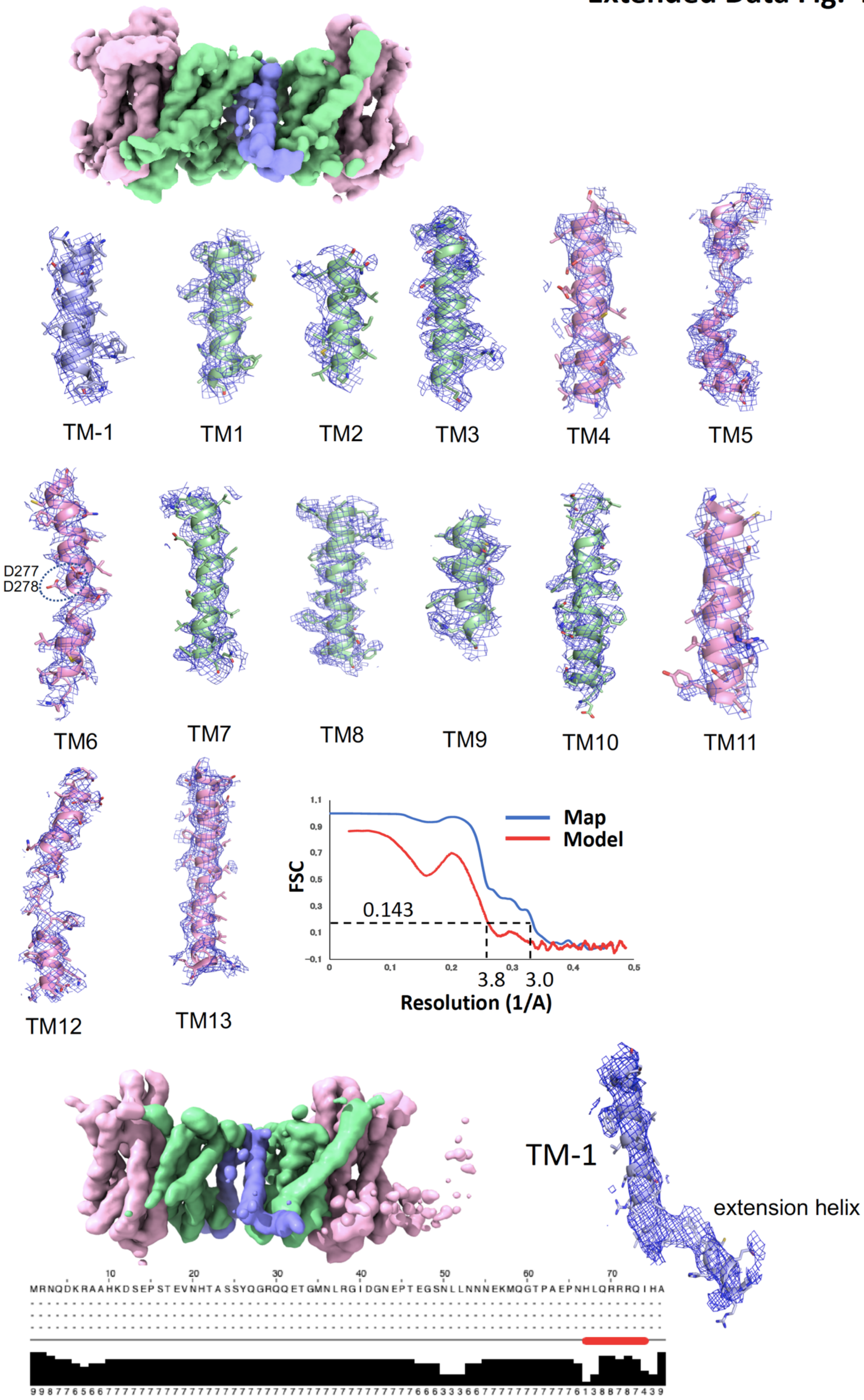

**Extended Data Fig. 4. Cryo-EM density of *bison* NHA2 $_{\Delta N}$ in detergent before and after density subtraction. a.** *above:* Cryo-EM density map of NHA2 $_{\Delta N}$ with the 6-TM core transport domains (coloured in pink), the dimer domain (coloured in green) and the N-terminal domain-swapped helix TM –1 (blue). *below:* cryo-EM density map and model are shown for all transmembrane segments for *bison* NHA2 $_{\Delta N}$ coloured is in the maps above residues D277 and D278 (encircled) on TM6 have been modelled after NapA at pH 8.0 (PDB id: 4bwz). The FSC curves for the *bison* NHA2 $_{\Delta N}$ in detergent after masking **b**. Cryo-EM density map of NHA2 $_{\Delta N}$ with the extension helix that could be modelled prior to masking and the corresponding FSC curve. The full-length *bison* NHA2 N-sequence was subjected to secondary structure prediction by Jpred (https://www.compbio.dundee.ac.uk/jpred/) with the predicted N-terminal helix shown as a red-bar.

#### Extended Data Fig. 5

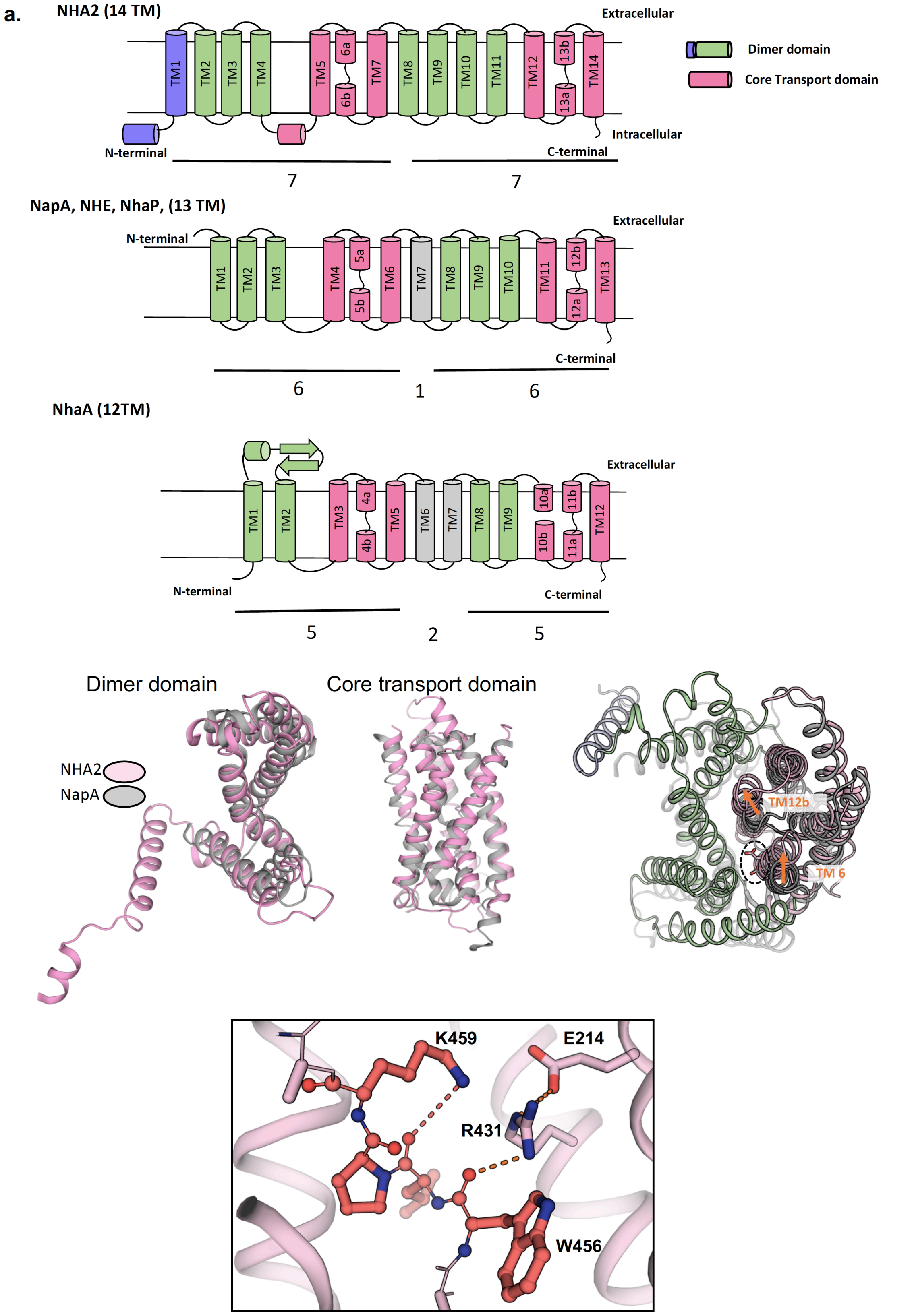

**Extended Data Fig. 5. Topology comparison of NHA2 to 13-TM and 12-TM $Na^+/H^+$ antiporter homologues, structural comparison to NapA, and the interactions in the NHA2 TM12a-b breakpoint. a.** *top:* NHA2 has 14 TMs with the N-and C-terminus located in the cytoplasm and is made up of two 7-TM structural-inverted repeats that form the 6-TM core domain (pink), dimerization domain (green) and N-terminal domain-swapped helix TM -1 (blue); note, the linker helix TM7 has been assigned with the dimer domain as is TM -1, even though technically both these helices could be assigned as independent from both core and dimer domains. *middle:* NapA, NHE and NhaP members with 13-TMs (N-terminus extracellular and C-terminus cytoplasm) is the most common topology seen in the $Na^+/H^+$ exchangers are made up of two 6-TM structural-inverted repeats that form the 6-TM core domain (pink), dimerization domain (green) and non-symmetry-related linker helix (grey). *bottom:* NhaA is the only Na+/H+ exchanger seen with 12-TMs is made up of two 5-TM structural-inverted repeats and forms the 6-TM core domain (pink), dimerization domain that contains a β-hairpin between TM1 and TM2 (green) and two non-symmetry related helices (grey). **b**. *left:* cartoon representation showing the structural superimposition of the respective domains of the NHA2 monomer, dimer domain and transport domain only (pink), against bacterial homologue NapA domains (grey). *right:* structural superimposition of the NHA2 (pink and green) and NapA (grey) monomers. The arrow highlights the structural differences in the core domain helices TM12b and TM6 between outward-facing NHA2 and NapA structures and the strictly conserved Asp278 (Asp157 in NapA) is shown in stick form. **c**. Cartoon representation of the ion-binding site of NHA2 with the TM12a-b breakpoint illustrated in stick form to highlight stabilization by K459 and R431, which is further salt-bridged to E214. Notably, these residues are likely to control the positioning of TM12b and its possible that the differences between the core domain in NapA and NHA2 shown in b., may reflect local, intermediate states.

#### Extended Data Fig. 6

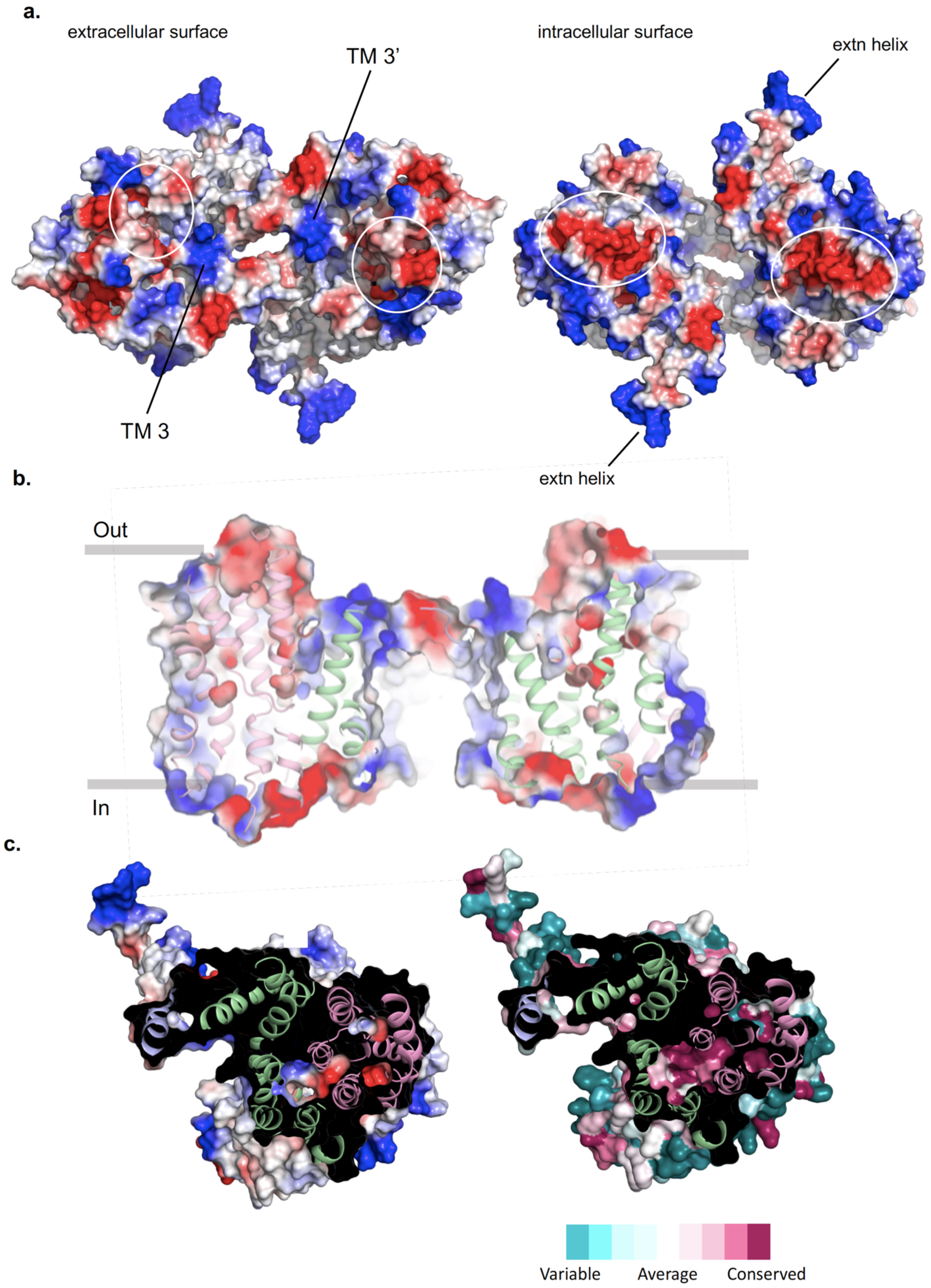

**Extended Data Fig. 6. Electrostatic surface potential *bison* NHA2$_{\Delta N}$ in detergent. a.** Electrostatic surface representation of the extracellular view of the outward-facing NHA2 homodimer (left) and the cytoplasmic view (right). The positive charges on the ends of TM3 and the N-terminal extension helix are highlighted. The circles show the positioning of the outward-facing funnel and location of the would be inward-facing funnel in the core domain. **b**. Electrostatic surface representation of the side view of the outward-facing NHA2 homodimer highlighting the large, intracellular gap between protomers. The oligomerization contacts on the extracellular side are mediated between TM-1 and TM8 on the neighbouring protomer. **c.** *left:* Cartoon representation of NHA2 from the extracellular side with the electrostatic surface representation through the ion-binding site of one monomer (coloured blue to red, for positive to negative charges, respectively). *right:* as in the left panel, but coloured according to conservation scores from the alignment of 500 mammalian NHA2 representative sequences calculated with ConSurf server[83].

### Extended Data Fig. 7

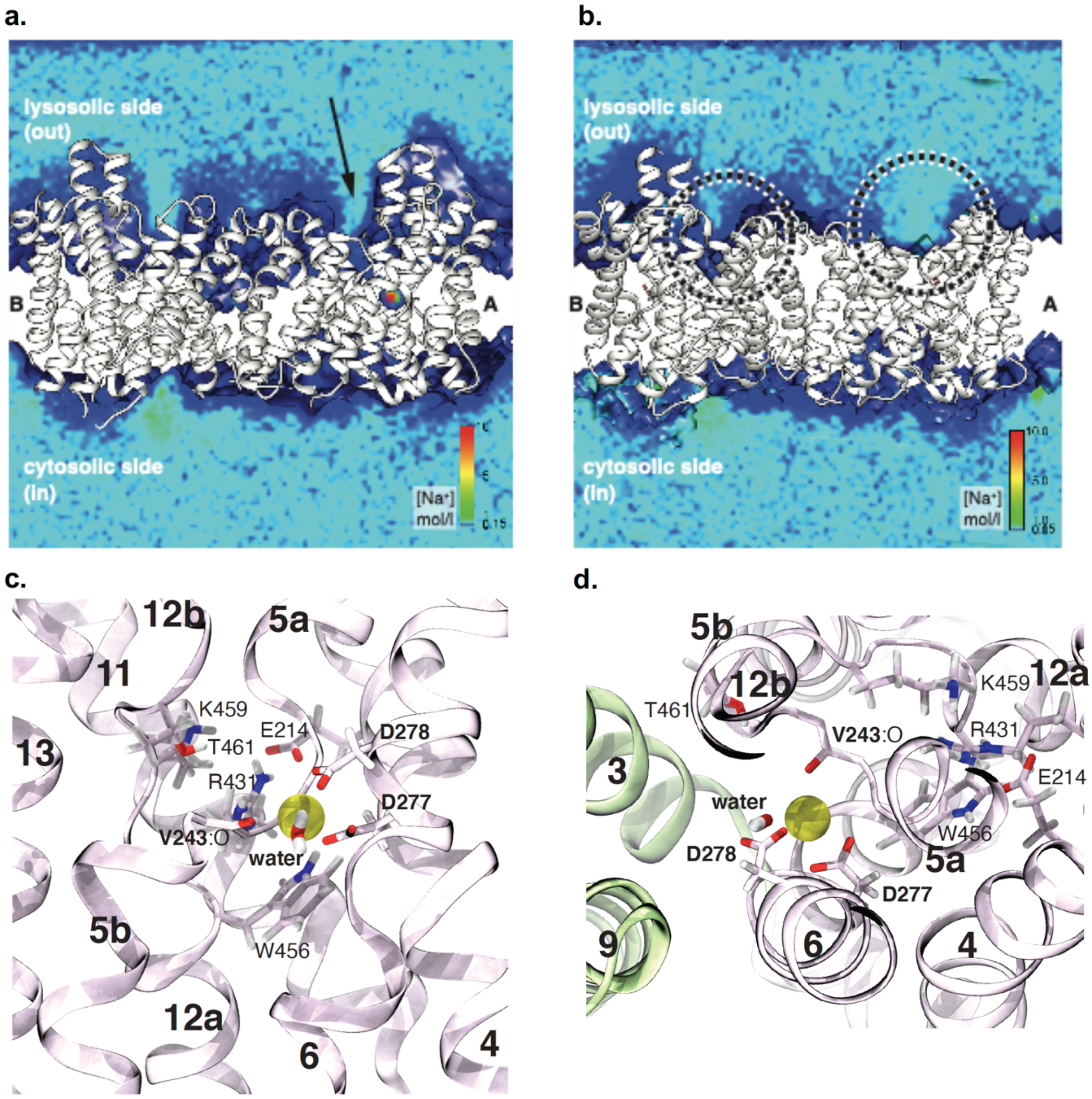


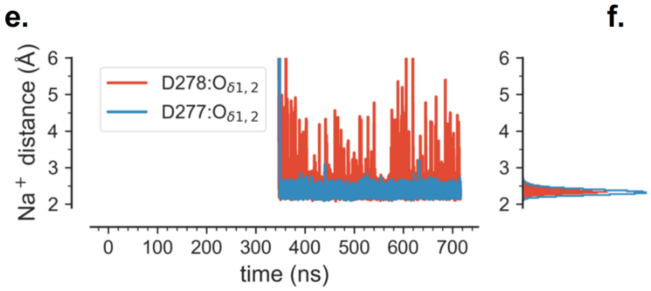


f.

| residue | n1 |
|---|---|
| Val243:O | 0.987 |
| Asp277-OD* | 1.992 |
| Asp278-OD* | 1.306 |
| water | 1.357 |
| **Total** | **5.642** |

**Extended Data Fig. 7. MD simulations of ion-binding to NHA2$_{\Delta N}$. a.** Density of sodium ions over 716 ns of simulation f-01-0 during which one sodium ion spontaneously entered the binding site in protomer A. The [NaCl] bulk concentration was 150 mM and ions were free to diffuse. The membrane was omitted for clarity. Protomer A (*right*) was simulated with both D277 and D278 deprotonated. Protomer B (*left*) had D277 protonated (neutral) while D278 remained negatively charged. The arrow indicates the outward facing entrance funnel through which ions diffuse to the binding site. **b.** Sodium ion density when either Asp278 is protonated and Asp277 is deprotonated (protomer A, *left*) or both aspartates are protonated (protomer B, *right*); based on simulation f-23-1. Ions do not enter the lysosolic funnel regions that are highlighted with dashed circles. **c**. Side view from the dimerization domain (omitted) on the putative sodium binding site, drawn from the last frame of the MD simulation. The sodium ion is shown in yellow, coordinating residues are indicated in bold, other residues near the binding site are labelled for context. Water molecules within 3 Å of the sodium ion were included, with only one present in this snapshot at the end of the MD trajectory. **d**. Top view (from the lysosolic side), with dimer domain in light green and core domain in light purple. **e**. Shortest distance of any sodium ion to either carboxylate oxygen in D277 or D278: timeseries (*left*) and histogram (*right*). **f**. Coordination of the bound sodium ion. The average contributions of oxygen atoms from different residues to the first hydration shell of the bound sodium ion, $n_1$, identify the binding site residues. "OD*" indicates the additive contributions from both the $O_{\delta 1}$ and $O_{\delta 2}$ carboxylate oxygen atoms; "water" from any water molecules. $n_1$<0.001 are not shown.

#### Extended Data Fig. 8

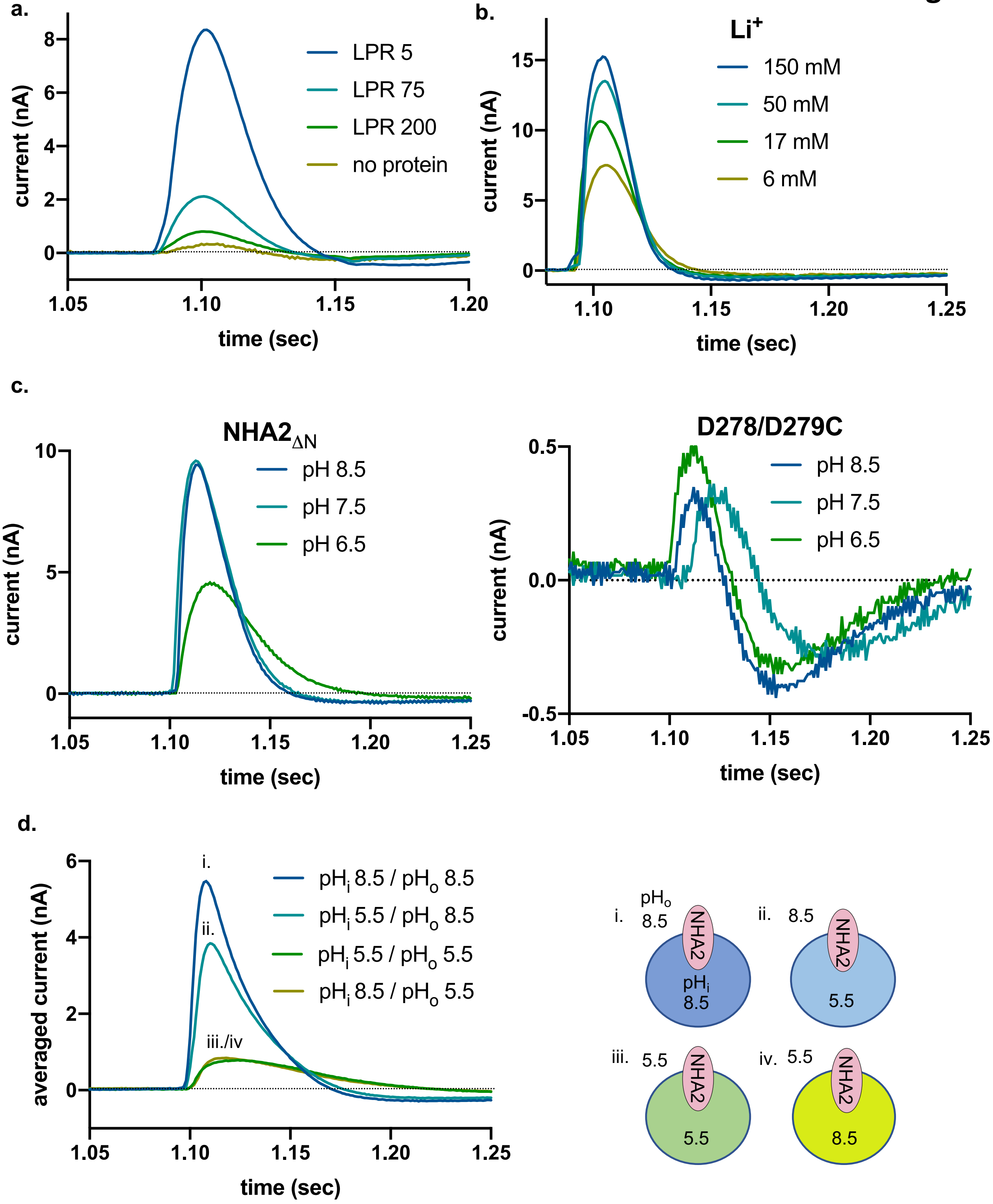

**Extended Data Fig. 8. SSM-based electrophysiology measurements of *bison* NHA2$_{\Delta N}$ proteoliposomes. a.** Transient currents in different lipid to protein ratios as indicated. **b.** Transient currents recorded after $Li^+$ concentration jumps at symmetrical pH 7.5. **c.** Transient currents recorded after addition of 150 mM NaCl at symmetrical pH 6.5, 7.5 and 8.5 for *bison* NHA2$_{\Delta N}$ (left) and *bison* NHA2$_{\Delta N}$ with Asp278 and Asp279 substituted with cysteine (right)**. d.** The pH was varied independently inside ($pH_i$) and outside ($pH_o$) the proteoliposomes, followed by 150 mM NaCl jumps to activate NHA2 in presence of pH gradient. Only lowering the pH on the outside dramatically affects current amplitudes, which is consistent with an electroneutral transport cycle[35,36]. Representative results of recordings performed on three individual sensors are shown.

### Extended Data Fig. 9

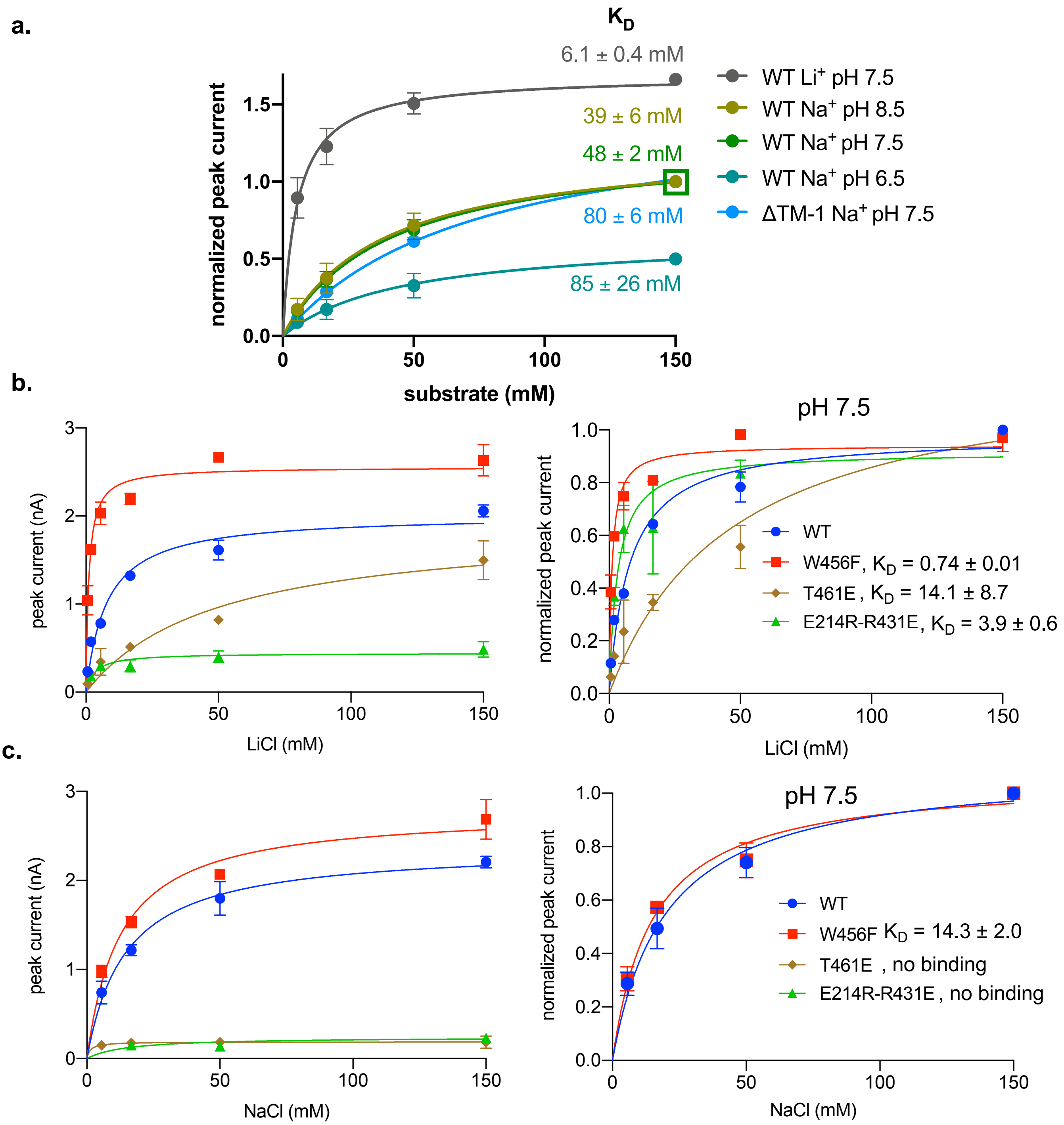

**Extended Data Fig. 9. SSM-based electrophysiology measurements of NHA2 proteoliposomes. a.** Fit of the normalized amplitude of the transient currents as a function of $Na^+$ ($Li^+$) concentrations and pH for *bison* NHA2$_{\Delta N}$ and *bison* NHA2$_{\Delta TM-1}$ and the corresponding binding affinity ($K_D$). Note, since we are measuring pre-steady-state ion translocation rather than steady-state currents, it is more accurate to refer this estimate as a binding constant ($K_D$) rather than the Michaelis–Menten constant, $K_{M,}$. Currents have been normalized for *bison* NHA2$_{\Delta N}$ 150 mM $Na^+$ at pH 7.5 as indicated by the green square **b**. The amplitudes of transient currents for *bison* NHA2$_{\Delta N}$ and derived *bison* NHA2$_{\Delta N}$ mutations recorded after $Li^+$ concentration jumps at symmetrical pH 7.5 (*left*) and the fit of the normalised amplitudes (*right*). **c**. As in b. for NaCl additions.

#### Extended Data Fig. 10

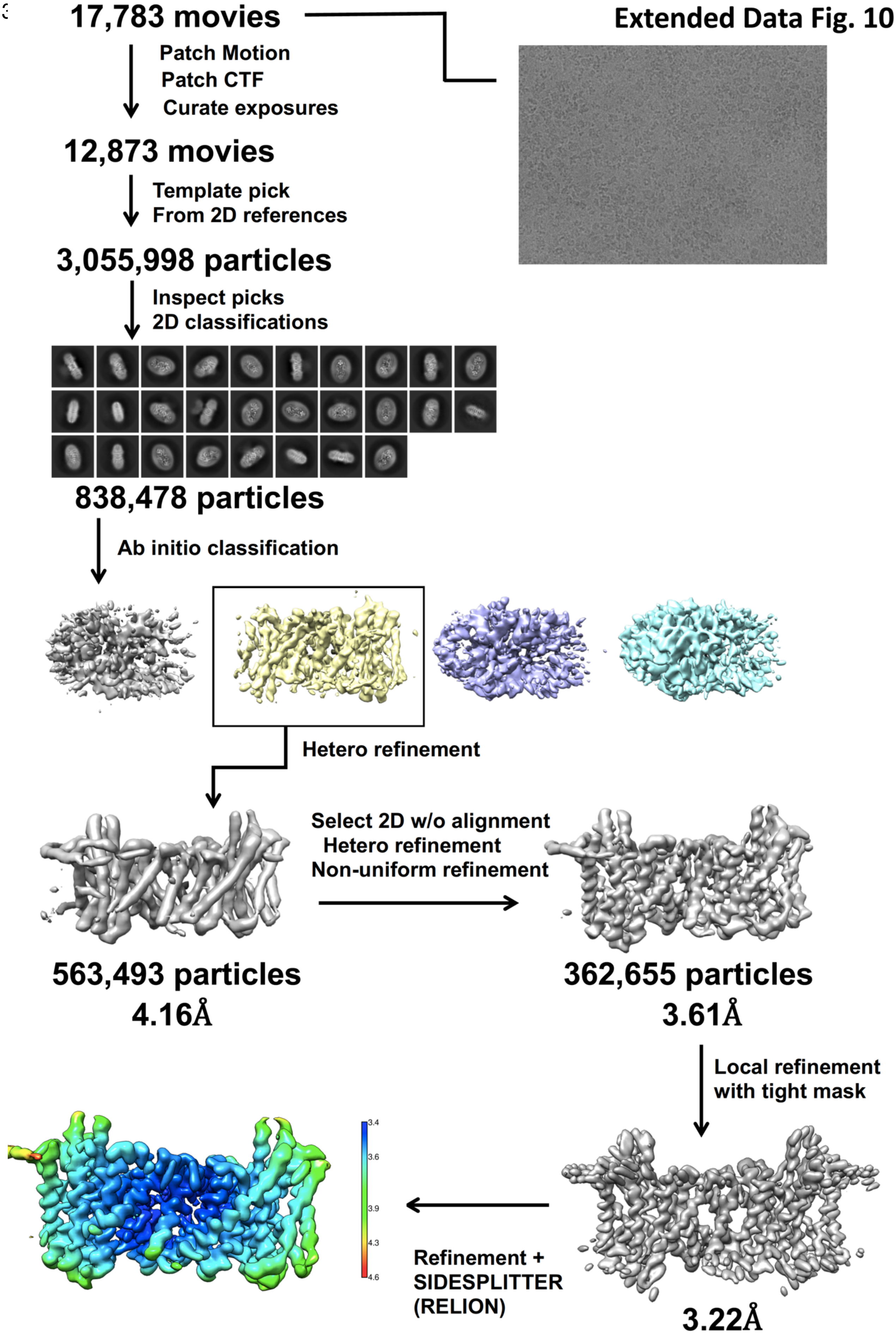

**Extended Data Fig. 10. The data-processing workflow of *bison* NHA2 $_{\Delta N}$ in nanodiscs.** The dataset contained 17,783 movies that were corrected by Patch motion correction and Patch CTF estimation in cryoSPARC. After template based autopicking, 3,055,998 particles were picked. Several rounds of 2D classification were performed, yielding 838,438 particles, which were subjected to 3D classification. One of the three 3D classes was selected, and it contained 563,493 particles. After non-uniform refinement in cryoSPARC, a resolution of 3.61Å was achieved that was imporved with local refinement and masking to 3.22Å and after SIDESPLITTER in Relion a resolution of 3.5 Å was achieved at gold standard FSC (0.143), with a local resolution range of 3.4 to 4.6 Å.

**Extended Data Fig. 11.**

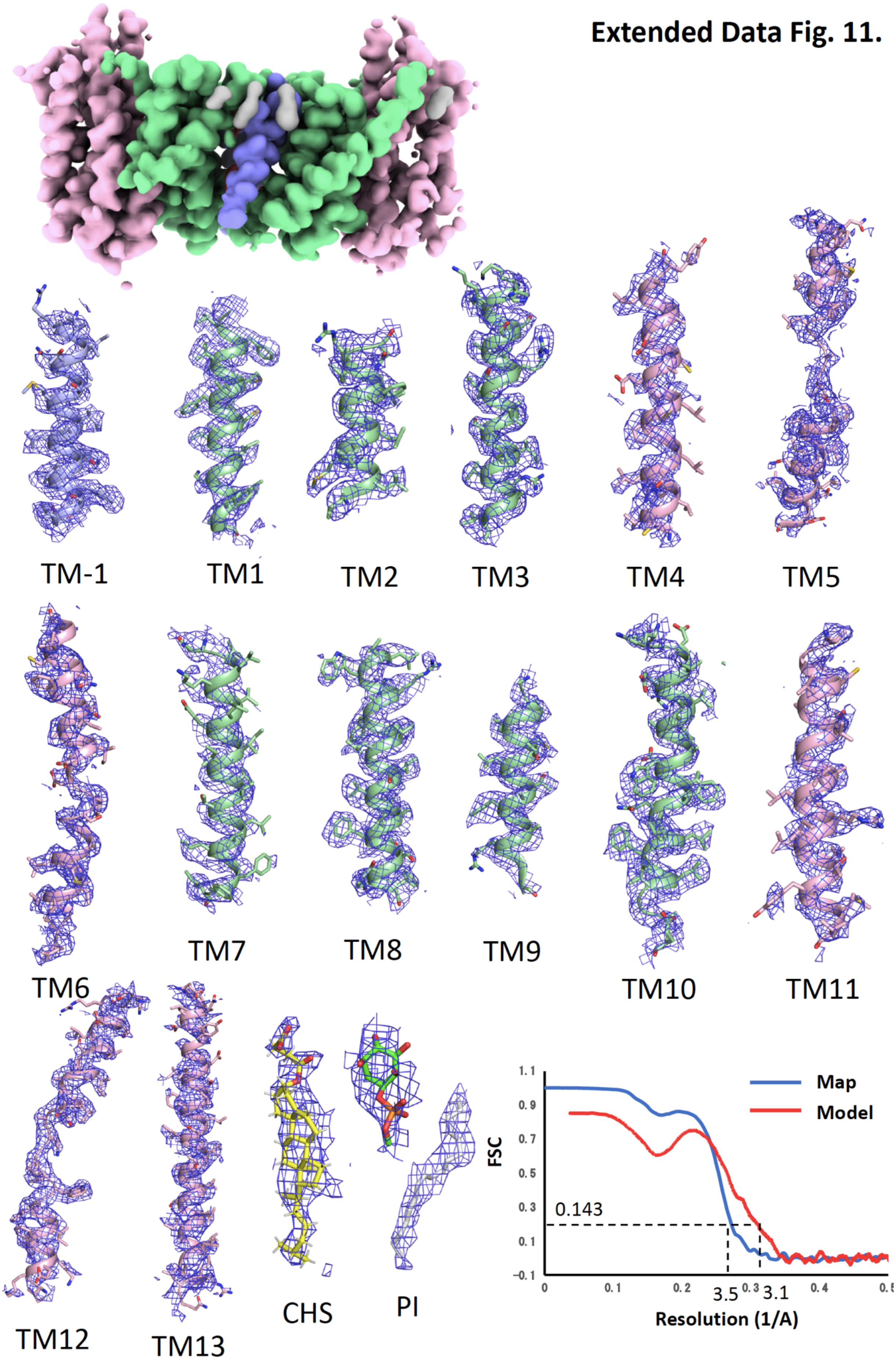

**Extended Data Fig. 11. Cryo-EM density of *bison* NHA2 $_{\Delta N}$ in nanodiscs.** Cryo-EM density map of NHA2 $_{\Delta N}$ in nanodiscs with the 6-TM core transport domains (coloured in pink), the dimer domain (coloured in green), the N-terminal domain-swapped helix TM –1 (blue) and cholesterol (grey). The cryo-EM density map and model are shown for all transmembrane segments for *bison* NHA2 $_{\Delta N}$ coloured is in the maps above. Representative cryo-EM density for the lipids is also shown. The FSC curves for the *bison* NHA2 $_{\Delta N}$ in nanodiscs after masking

#### Extended Data Fig. 12.

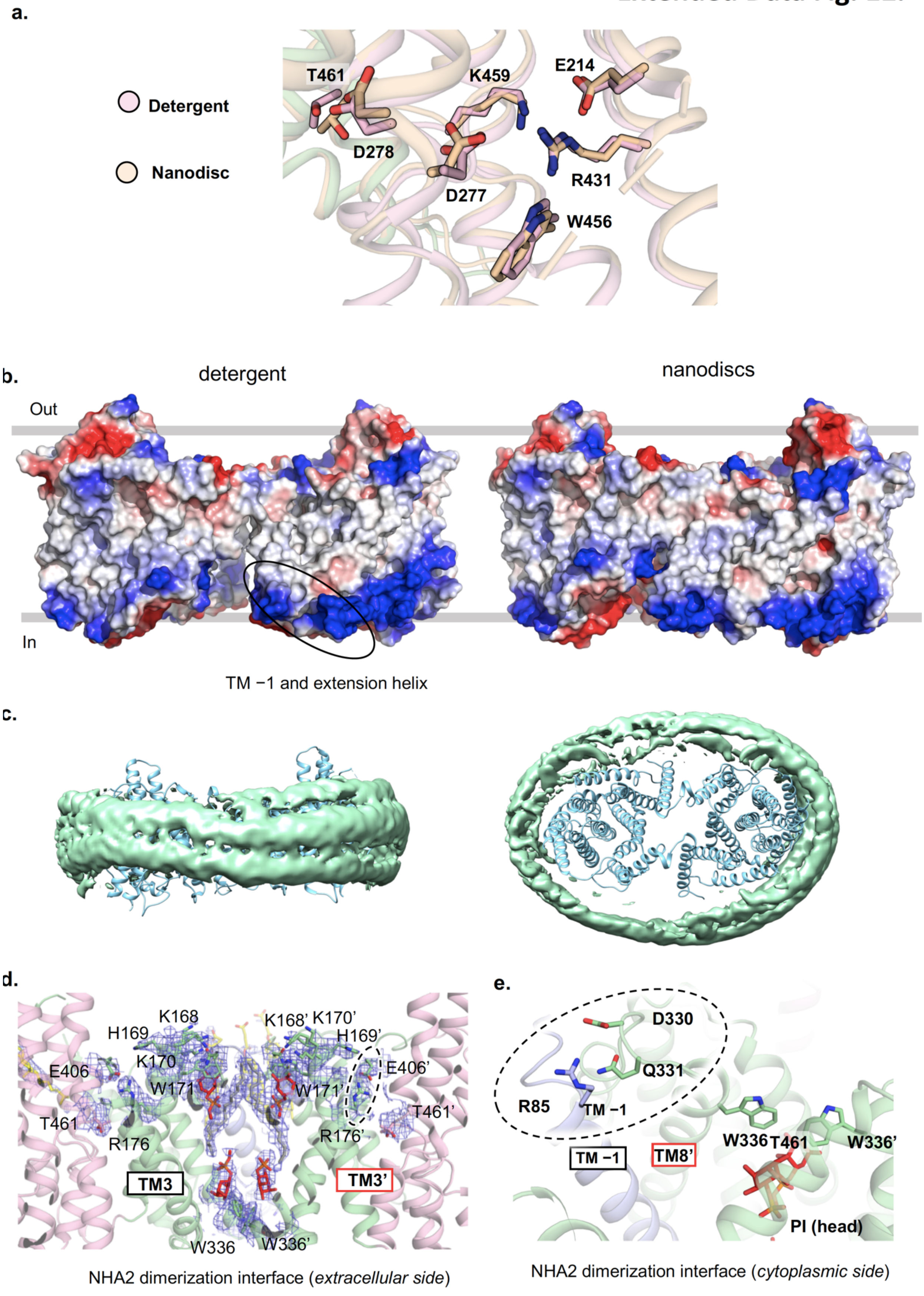

**Extended Data Fig. 12. Comparison of *bison* NHA2$_{\Delta N}$ structures in detergent and nanodiscs. a.** Comparison between ion-binding site residues in the NHA2$_{\Delta N}$ structure obtained in detergent and nanodiscs. **b**. Electrostatic surface representation of the side view of the outward-facing NHA2 homodimer highlighting the large, intracellular gap between protomers and intracellular positively-charged surface in detergent (*left*) that is closed upon TM −1 rearrangement in nanodiscs incorporated with PI lipids (*right*). **c**. The cyo EM density for the nanodisc from the side and top and the manually placement of the *bison* NHA2$_{\Delta N}$ structure in detergent. **d**. Cartoon representation on the extracellular side showing the polar contacts between the K168, H169, K170 and W171 residues in the end of TM3 of one protomer (green sticks, labelled) with PI lipids located between the same interactions in the other protomer in nanodiscs. (green sticks, labelled with ‘). Connected to TM3 is R176 that interacts with E406 (TM10) that might stabilise the outward-facing cavity to the ion-binding site as highlighted with T461 in the core domain (pink). PI lipids (red sticks), cholesterol located on extracellular half of the protein (yellow sticks). **e.** Cartoon representation on the cytoplasmic side showing the polar contacts between strictly-conserved D330, Q331 residues in the TM8-TM9 loop of one protomer (green sticks, labelled) and Arg85 in TM −1 of the other protomer in nanodiscs (blue sticks, labelled) wherein TM8 moves inwards to coordinate PI lipids (red sticks) via lipid-interface tryptophan residues that are highly conserved (see ED Fig. 1).

### Extended Data Fig. 13.

**a.**

Intrinsic TM-1 dynamics of NHA2: lowest to mid energy modes

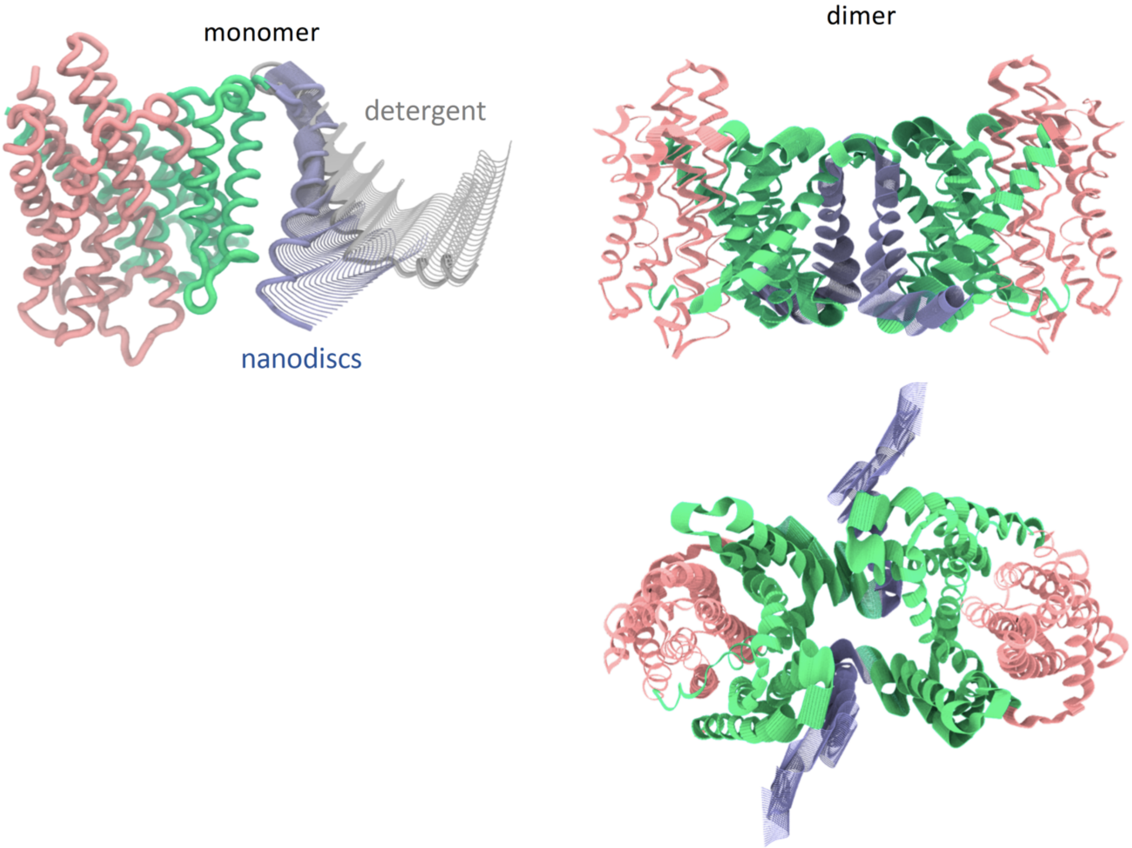


**b.**

Intrinsic elevator-like motions of NHA2: mid-energy modes

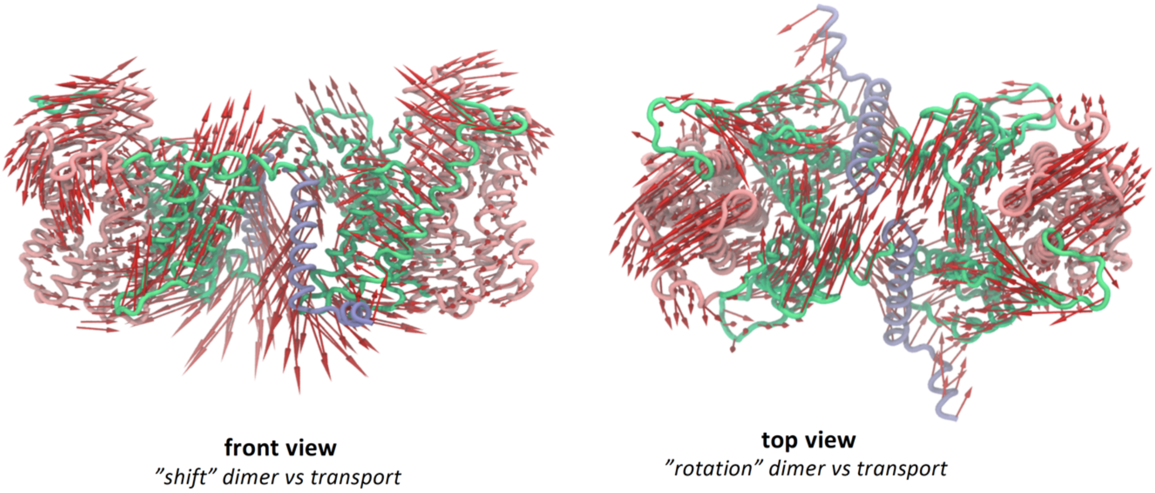

**Extended Data Fig. 13. Elastic Network Modelling of NHA2 dimers and monomers.** **a.** Intrinsic TM −1 dynamics of both monomeric (left) and dimeric (right) NHA2. The lowest frequency modes of bison NHA2$_{\Delta N}$ are dominated by intrinsic TM−1 dynamics that spontaneously interconvert to the experimental conformations seen between detergent and nanodisc structures. **b**. Elevator-like motions of both monomeric and dimeric NHA2. Mid-range modes (NM12-18) encode displacements of the dimer versus the transport domain that can drive transitions to inward-like states resembling NapA inward conformation.

#### Supplementary Fig. 1

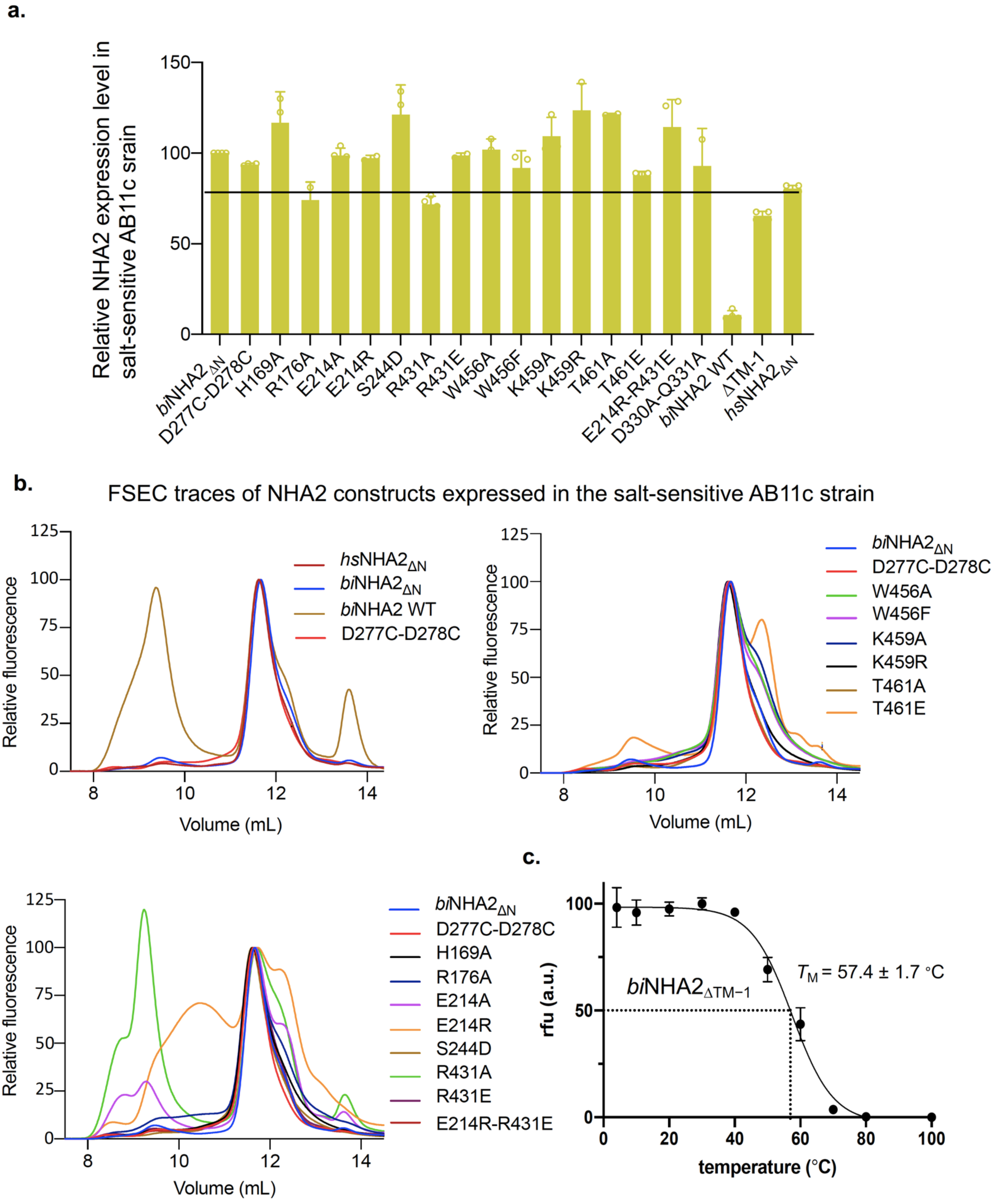

**Supplementary Data Fig. 1. Expression and quality of NHA2-GFP fusions assessed by fluorescence-detection size exclusion chromatography (FSEC) in the ABc11 yeast strain**. **a**. Bar represent normalized expression level compared to *bison* NHA2 $_{\Delta N}$ for various NHA2 constructs and mutants as measured by GFP fluorescence in yeast whole-cells. In all experiments errors bars represent the range of n = 2 to 4 independent cultures; note, only the full-length bison NHA2 construct is poorly expressed (10-fold less than *bison* NHA2 $_{\Delta N}$) and has a large fraction of aggregates (see b.), yet still shows significant complementation at 10 mM LiCl and some complementation at 25 mM LiCl (Supplementary Fig. 3a). **b.** FSEC traces of listed NHA2 constructs extracted by DDM/CHS from Abc11 yeast membranes. The peak at 11.5 ml corresponds to the NHA2 homodimer and all the constructs shown here have a main FSEC peak at 11.5ml; the only constructs with a shifted predominant peak at 12.2 ml for the monomer are NHA2 $_{TM-1}$ and NHA2 $_{\Delta N}$ (Q330A, D331A) shown in Fig. 3b. All constructs showing a minor monomer peak at 12.2 ml show poor folding as apparent by the aggregation peak in the void. e.g., E214R, R431A, T461E, and *bison* NHA2 full-length (WT). **c**. Thermal shift of purified monomeric NHA2 $_{TM-1}$-GFP in the presence of DDM/CHS. Data presented are normalized mean fluorescence as mean values ± data range of n = 3 technical repeats; the apparent $T_M$ was calculated with a sigmoidal 4-parameter logistic regression function; the average $\Delta T_M$ presented is calculated from n = 2 independent titrations.

#### Supplementary Fig. 2

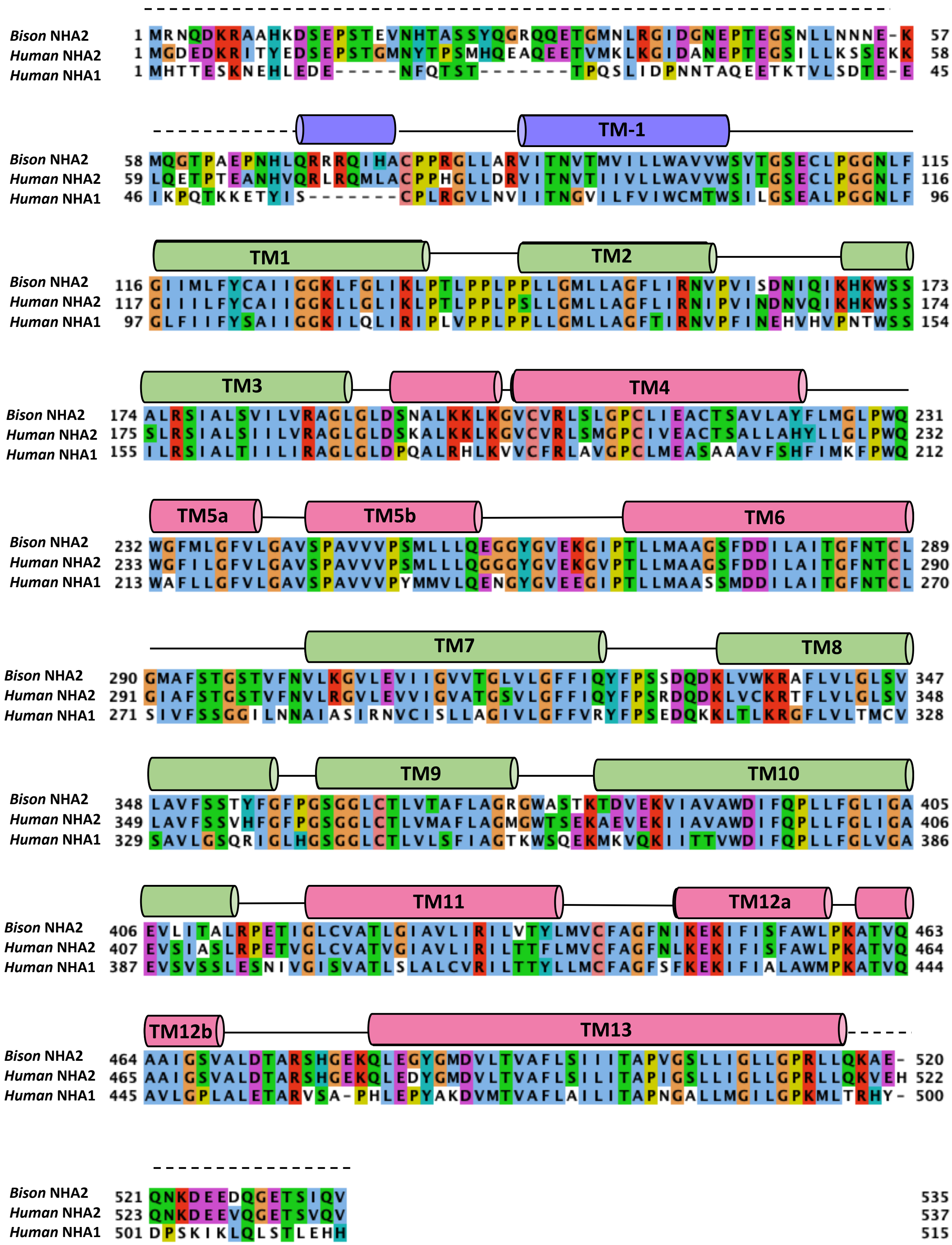

**Supplementary Fig. 2 Multiple sequence alignment of *bison* NHA2 and *human* NHA1 and NHA2 sequences.** NHA2 sequences were aligned by clustal omega and coloured by residue type in Jalview. Breakpoints (s-shaped line), core 6-TM transport domain TMs (pink), dimerization domain (green) and domain-swapped helix TM –1 (blue) are indicated.

#### Supplementary Fig. 3

##### LiCl sensitivity

**a.**

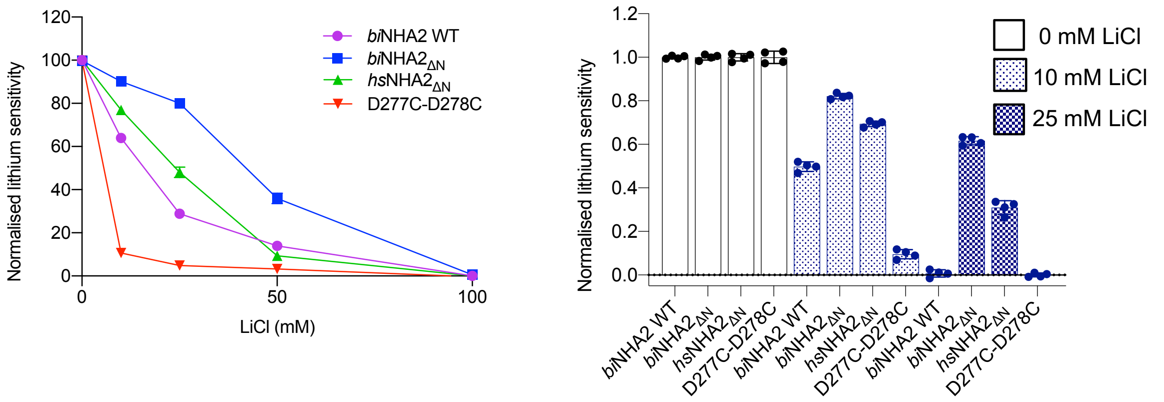


**b.**

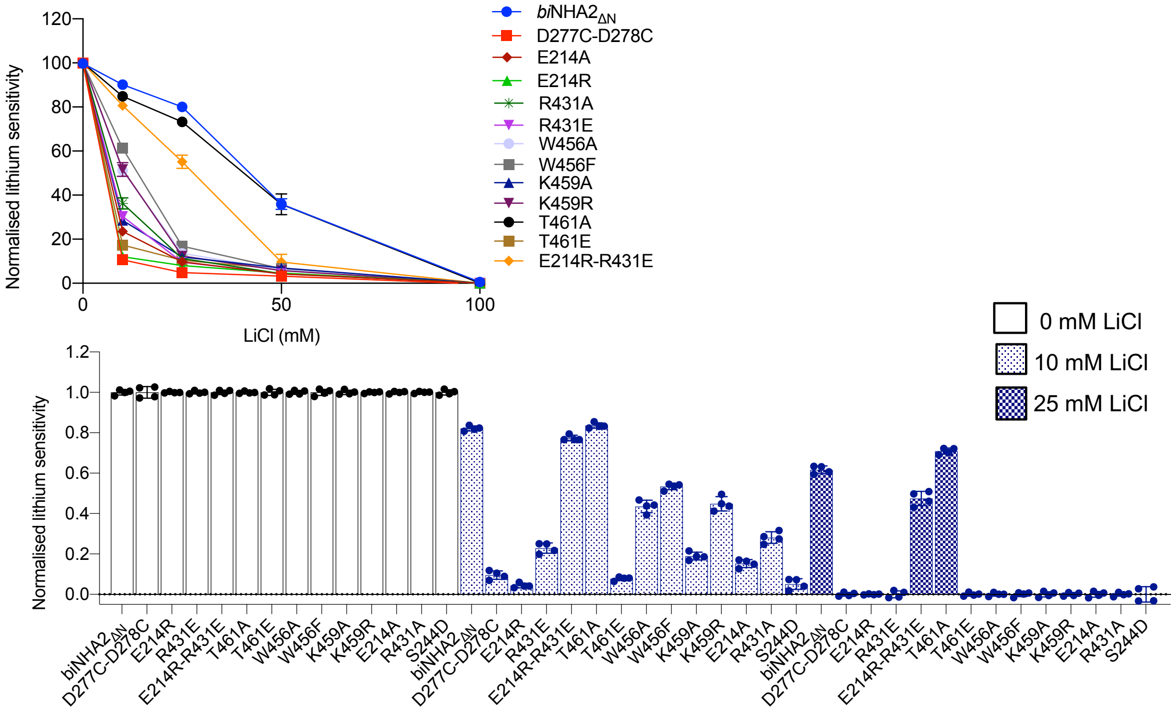


**c.**

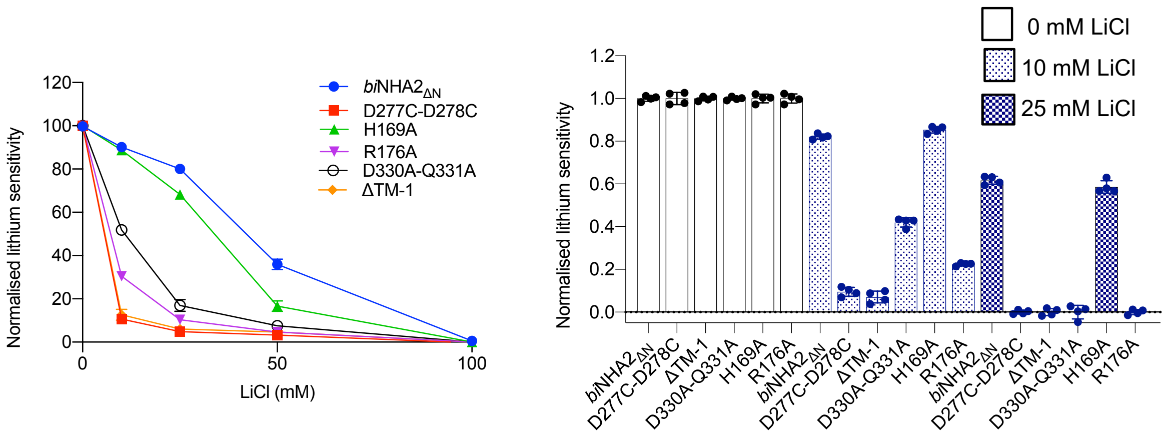

**Supplementary Fig. 3. Functional complementation of $Li^+$ sensitivity in the yeast strain AB11c by heterologous expression of NHA2 constructs**. **a**. *above:* Ab11c strain was transformed with *bison* NHA2 WT, *bison* NHA2 $_{\Delta N}$, *human* NHA2 $_{\Delta N}$ or *bison* NHA2 $_{\Delta N}$ with both Asp277 and Asp278 have been substituted with cysteine (D277C-D278C); *human* NHA2 numbering is D278 and D279. Yeast were grown as outlined in Methods in -URA media supplemented with 2% galactose and LiCl and growth was determined by optical density of the culture at 600 nm ($OD_{600}$) after 48 at 30°C and normalised against growth of *bison* NHA2 $_{\Delta N}$. *below:* To facilitate comparison the above data is shown as bar graphs. In all experiments described the errors bars, s.e.m.; n = 4 independent cultures. **b**, As shown in a., for ion-binding site mutations of *bison* NHA2 $_{\Delta N}$, which were normalised against growth of *bison* NHA2 $_{\Delta N}$ that was always cultured in parallel **c**. as shown in a. for structural mutants of *bison* NHA2 $_{\Delta N}$ .

**Supplementary Fig. 4**

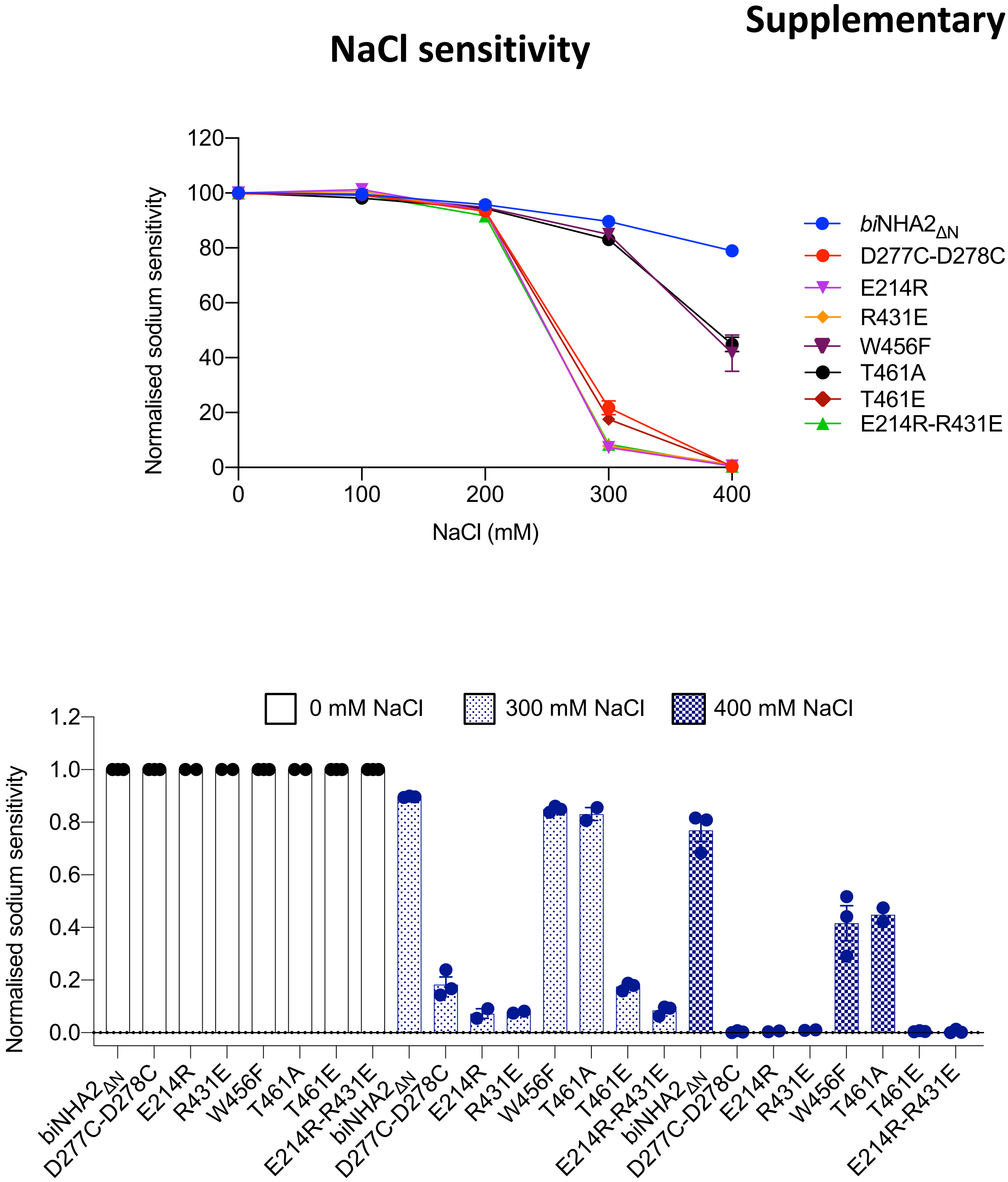


**Supplementary Fig. 4. Functional complementation of Na+ sensitivity in the yeast strain AB11c by heterologous expression of *bison* NHA2 $_{\Delta N}$ ion-binding site mutants.** In most cases Li+ was used to asses NHA2 complementation as Li$^{+ i}$s more toxic to yeast cells and NHA2 has a higher affinity ($K_D$) for Li$^{+}$ vs. Na$^{+}$. Nevertheless, for those ion-binding site mutants that retained some Li$^{+}$-complementation they were further assessed for $Na^+$ sensitivity. *above:* Yeast were grown as outlined in Methods in -URA media supplemented with 2% galactose and NaCl and growth was determined by optical density of the culture at 600 nm ($OD_{600}$) after 72 at 30°C and normalised against growth of *bison* NHA2 $_{\Delta N}$. In all experiments described the n = 2 or 3 independent cultures. *below:* To facilitate comparison the above data is shown as bar graphs.

#### Supplementary Fig. 5

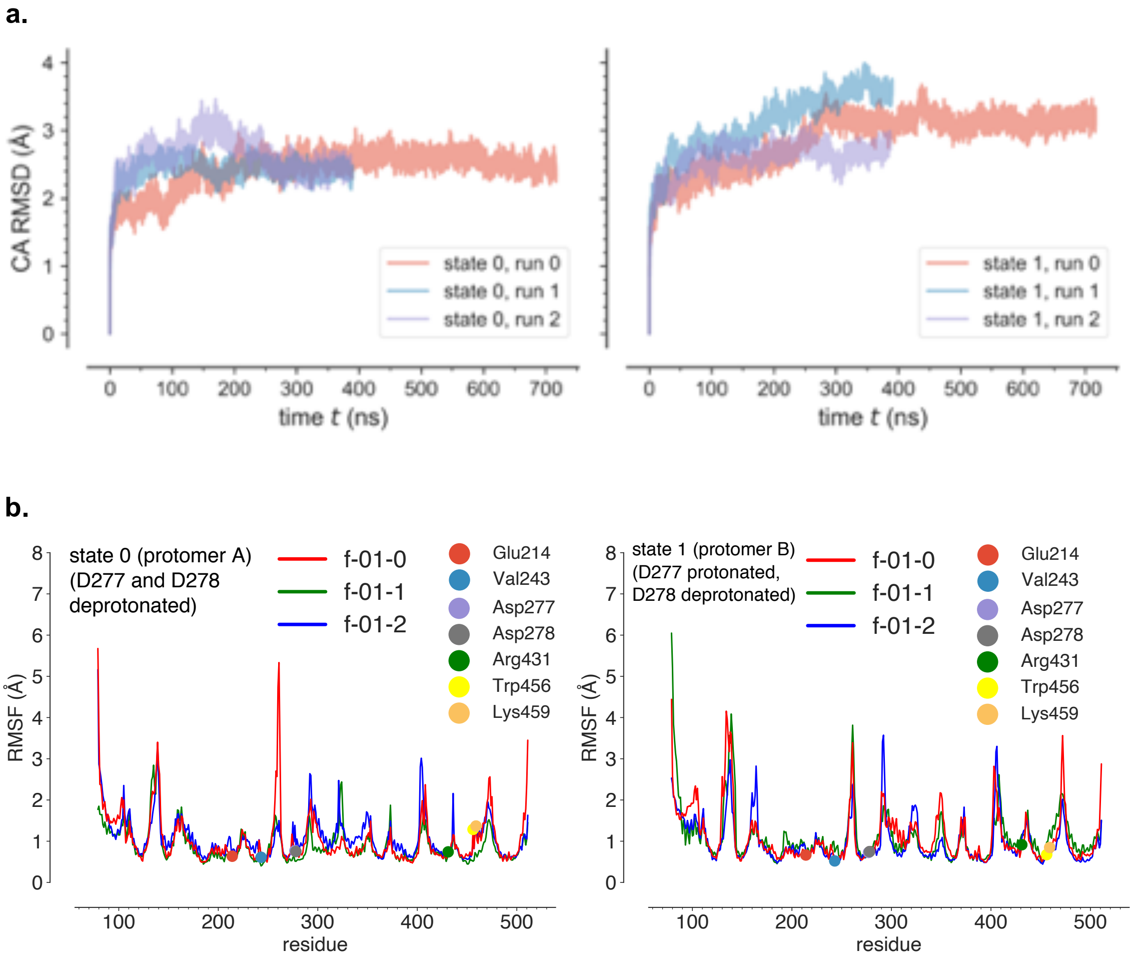


**Supplementary Fig 5. Average structural variability in MD simulations.** Conformational drift from the initial experimental structure and local flexibility was assessed with root mean square quantities. Protomers A and B, which were simulated in different protonation states were structurally superimposed on themselves and analyzed independently. *Left:* state 0 (D277 and D278 deprotonated). *right:* state 1 (D277 protonated, D278 deprotonated). **a.** $C_\alpha$ root mean square distances (RMSD) from the first trajectory frame after energy minimization and short equilibration MD. **b.** $C_\alpha$ root mean square fluctuations (RMSF) Locations of key residues are highlighted as colored circles.

#### Supplementary Fig. 6

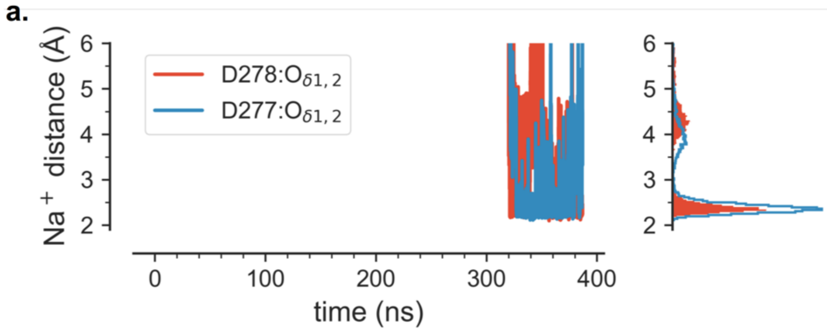


b.

| residue | n1 |
|---|---|
| Gly274:O | 0.361 |
| Ser275:O | 0.035 |
| Asp277-OD* | 1.343 |
| Asp278-OD* | 0.956 |
| Gly360:O | 0.001 |
| water | 3.491 |
| **Total** | **6.187** |

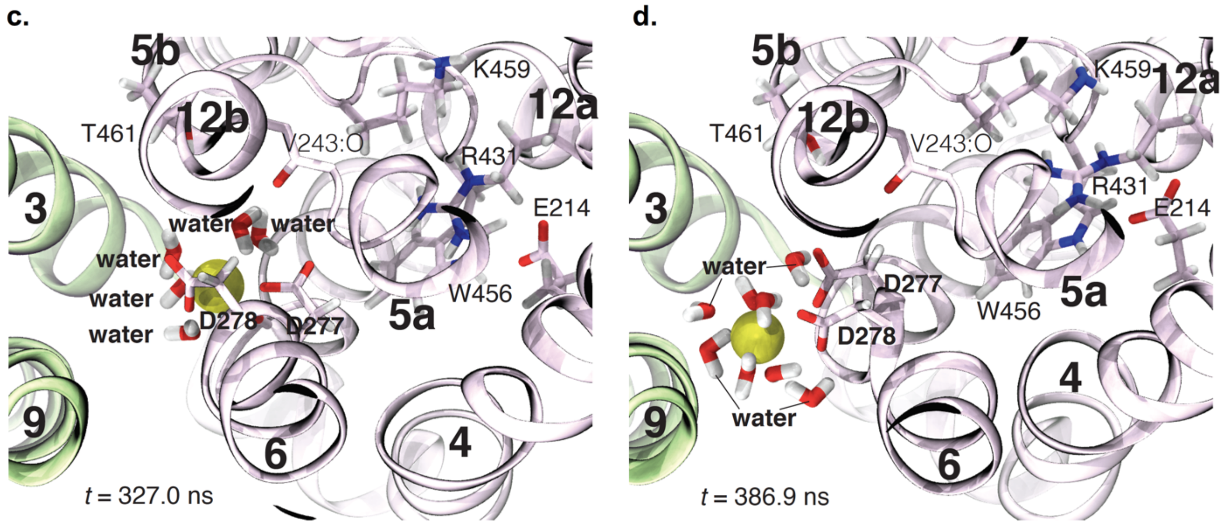


**Supplementary Fig 6. Initial ion binding event in MD simulations**. In simulation f-01-2, a sodium ion partially bound to the ion binding residues in protomer A (protonation state 0, i.e., D277 and D278 negatively charged). **a.** Shortest distance of any sodium ion to either carboxylate oxygen in D277 or D278: timeseries (left) and histogram (right). **b.** Coordination of the partially bound sodium ion, quantified by the average contributions of oxygen atoms from different residues to the first hydration shell of the bound sodium ion, $n_1$. $n_1$<0.001 are not shown. Only backbone carbonyl oxygen atoms ("O") or the delta oxygen atoms of the carboxylate groups participated in binding. **c**. Top view (from the lysosolic side), with dimer domain in light green and core domain in light purple at time 327.0 ns. Water molecules within 3 Å of the sodium ion (yellow) are included. **d**. Top view at the end of the simulation at 386.9 ns.

**Supplementary Video 1. Morph showing the structural transitions between the detergent and nanodisc *bison* NHA2 $_{\Delta N}$.. a**. NHA2 structural transitions as viewed from the extracellular side with core domain (pink), dimerization domain (green) and domain-swapped TM −1 (blue). **b**. As in a., as viewed from the side. **c**. As in a., as viewed from the cytoplasm.

**Supplementary Video 2. Intrinsic dynamics of TM −1 in *bison* NHA2 $_{\Delta N}$. a.** Intrinsic dynamics of the NHA2 monomeric in nanodiscs with core domain (pink), dimerization domain (green) and domain-swapped TM −1 (blue) that spontaneously forms a conformation like that seen in detergent with TM −1 (grey). **b**.
